# Integrated transcriptomics and proteomics define the TRP channel hierarchy in mouse cortex

**DOI:** 10.64898/2026.04.07.716663

**Authors:** Muhammad Bilal, Karthik Subramanian Krishnan, A.J. Sethi, Natasha Vassileff, Jereme G. Spiers, Rippei Hayashi, Ehsan Kheradpezhouh

## Abstract

Transient receptor potential (TRP) channels are evolutionarily conserved polymodal cation channels that mediate diverse sensory functions across the animal kingdom. Although TRP channels play key roles in peripheral sensation, their expression and functional relevance in the cerebral cortex remain poorly defined. Here, we integrate long- and short-read transcriptomics, targeted qPCR and membrane-aware proteomics to quantify TRP family members in adult mouse cortex. Across transcriptomic platforms, cortical TRP expression is dominated by TRPML, TRPC, and TRPM subfamilies, with lower representation of TRPP/TRPV, whereas *Trpa1* and *Trpv1* lie near empirical detection thresholds. Our proteomic workflow yields reproducible protein-level evidence for a subset of cortical TRPs, including TRPV2, TRPC4, TRPM3, TRPM7 and TRPP2, consistent with transcript rank order, while TRPA1/TRPV1 do not meet replicate-level protein-group detection criteria under 1% FDR control. Together, these multi-platform measurements establish a quantitative reference for cortical TRP biology and a framework for profiling low-abundance ion channels in complex brain tissue.

## Introduction

To navigate and survive in dynamic environments, animals evolved sophisticated sensory systems that convert external stimuli into neuronal activity. A major molecular component is the Transient Receptor Potential (TRP) ion channel superfamily, which is evolutionarily conserved across eukaryotes and comprises 28 paralogous members in mammals^1,2^. This diversification enabled functional specialisation, with TRP channels mediating detection of thermal, chemical, and mechanical cues^3–5^ and modulating membrane excitability through calcium (Ca²⁺) influx^6,7^. Given that their functions are traditionally associated with the peripheral nervous system, TRP-dependent activity in the cortex, where cells are insulated from direct environmental exposure and primarily integrate network signals, implies intrinsic modulatory roles within central circuits^2,7,8^. TRP channels have been proposed to tune tactile and visual sensitivity according to context, arousal and expectation^2,9,10^, and are implicated in disorders characterised by Ca²⁺-dependent network hyperexcitability^3,8,11^. However, their cortical expression patterns and the molecular mechanisms underlying such modulation remain poorly defined, particularly whether TRP channels exhibit robust basal expression or are conditionally engaged under specific physiological or pathological states.

TRP channels comprise seven subfamilies: TRPC (canonical), TRPV (vanilloid), TRPM (melastatin), TRPA (ankyrin), TRPML (mucolipin), TRPP (polycystin), and TRPN (NOMPC-like, absent in mammals). Among these, Group I members (TRPC, TRPV, TRPM, and TRPA) are the most functionally implicated in neuronal excitability, Ca^2+^ homeostasis and synaptic plasticity^3,12,13^. Within this group, seminal work on archetypal channels TRPA1 and TRPV1 established their roles as polymodal sensors in peripheral nociception, thermosensation, and mechanotransduction^14–17^. Given their robust expression in dorsal root ganglia (DRG) and well-defined sensory functions^15,18–20^, both channels have been proposed as compelling candidates to influence synaptic and network activity in the brain^21–23^. However, resolving their cortical expression has been hindered less by biological disagreement than by technical constraints in quantifying low-abundance, multi-pass membrane proteins^24–27^. In long-lived neurons, temporal regulation of mRNA transcription, the transient nature of mRNA translation, and slow protein turnover lead to discordance between steady-state mRNA and protein abundances^28–30^. Conventional Liquid Chromatography – Tandem Mass Spectrometry (LC–MS/MS) exacerbates these constraints because TRP channels exhibit pronounced hydrophobicity, assemble into large multimeric architecture with limited solubility, and yield few proteolytic peptides, resulting in systematic underrepresentation^31–35^.

Together, these uncertainties highlight the absence of a coherent transcript- and protein-level account for TRP channel expression in the cerebral cortex, limiting our ability to define how TRP-mediated Ca²⁺ signalling contributes to cortical computation, plasticity, and disease. Addressing this gap requires an approach that resolves TRP transcript diversity while providing orthogonal protein-level evidence for low-abundance, hydrophobic multi-pass channels. Here, we combine long- and short-read transcriptomics, targeted qPCR, and membrane-aware proteomics to generate a map of TRP channel expression at a basal physiological level in the adult mouse cortex. Beyond detection, this integrated approach defines empirical limits and quantitative upper bounds for channels that remain below robust biochemical detectability. This work establishes a reference baseline for systematic investigation of context-dependent TRP regulation across plasticity, neuroinflammation and neurological disease.

## Results

### Integrated RNA-seq defines the TRP transcript landscape in adult mouse cortex and dorsal root ganglia

To quantify TRP transcript abundance in adult mouse cortex while retaining full-length isoform resolution, we performed Nanopore direct RNA sequencing across three biological replicates (workflow shown in Fig. 1A and Supplementary Fig. 1a). Out of 28 mammalian TRP genes, we identified transcripts of 22 genes, spanning all major subfamilies; most were detected at low transcript per million (TPM; Fig. 1B; Supplementary Fig. 3a and Supplementary Table 1). The most abundant TRP channels (in mean TPM) were *Mcoln1* (TRPML1; ∼57), *Trpc1* (∼34), *Trpm2* (∼17), *Trpv2* (∼17), *Trpm7* (∼16) and *Pkd2* (∼15). Intermediate expression was observed for *Trpm3* (∼10) and *Trpc4*, *Trpc6* and *Trpm4* (∼7 each). In contrast, *Pkd2l2* (TRPP5), *Trpa1*, *Trpc2*, *Trpc5*, *Trpm1*, *Trpm6*, *Trpv1*, *Trpv3* and *Trpv6* were near background (≤1 TPM), consistent with negligible basal transcription in adult cortex. *Mcoln2*, *Mcoln3*, *Pkd2l1*, *Trpm5*, *Trpm8* and *Trpv5* were not detected across replicates (falling below conservative RNA-seq detection thresholds), suggesting highly restricted or absent cortical expression under baseline conditions. Read count-based visualisations of the same Nanopore libraries (Supplementary Fig. 2a) recapitulated the TPM-derived rank order and further illustrated the sensitivity of long-read sequencing for capturing low-abundance full-length transcripts. In addition, because TRP genes can exhibit isoform diversity that is difficult to resolve with short-read sequencing, we next used NanoCount for reference-based transcript quantification and IsoQuant to reconstruct TRP isoform structures from Nanopore reads. This analysis identified seven candidate TRP transcript models that were not represented as complete transcript models in Ensembl release 115 (Supplementary Figs. 4a,b; a representative example in *Trpv2*, Supplementary Fig. 5; Supplementary Table 2). These candidate isoforms were reproducibly detected at low abundance and were mainly associated with alternative splice-site usage and/or exon skipping, highlighting them as candidates for future functional validation in the mammalian brain. Nanopore direct RNA sequencing enabled the prediction of chemical modification on TRP mRNAs. We identified N6-methyladenosine (m^6^A)-associated signal patterns at varying levels across TRP transcripts, and they are mostly found in DRACH (D = A/G/U, R = A/G, H = A/C/U) or RAC sequence motifs, consistent with canonical m⁶A deposition sites (Supplementary Fig. 6).

**Fig. 1:**
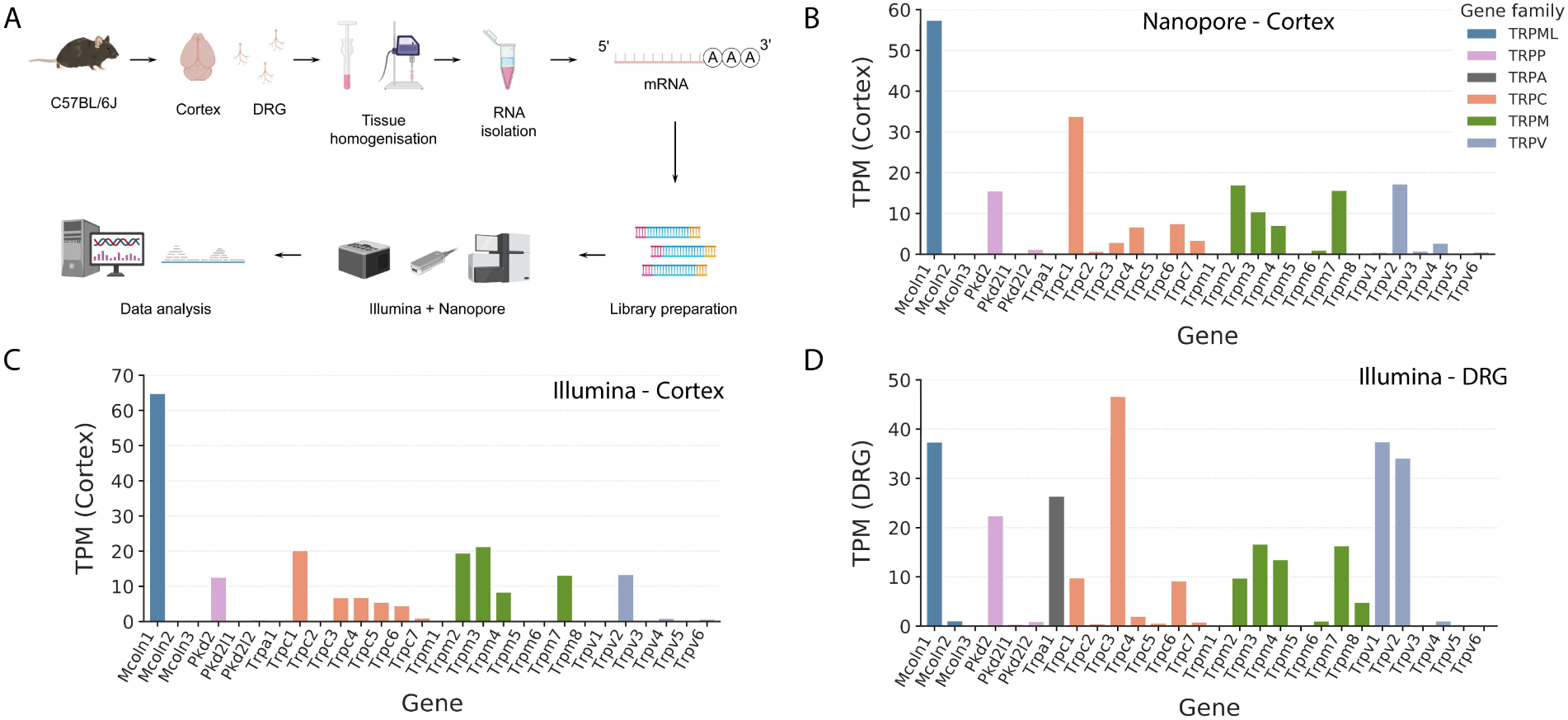
Integrated RNA-seq profiles of TRP channels in mouse cortex and dorsal root ganglia (DRG). (**A**) Experimental workflow for transcriptomic analysis. Cortical poly(A)+ RNA was used for Nanopore direct RNA sequencing, whereas Illumina short-read RNA-seq was performed on poly(A)+ enrichment RNA from cortex and dorsal root ganglia (DRG). Libraries were prepared separately and sequenced together in a multiplexed Illumina run. (**B**) Nanopore direct RNA-seq (mean transcripts per million (TPM); *n = 3* biological replicates) of cortical TRP transcripts; bars are coloured by TRP subfamily. (**C**) Illumina RNA-seq quantification of cortical TRP transcripts (mean TPM, *n = 3* biological replicates). (**D**) Illumina RNA-seq quantification of TRP transcripts from pooled DRG (3 mice; *n = 1* pooled library). Genes are shown in a consistent order across panels.

Illumina short-read RNA-seq (analysis pipeline, Supplementary Fig. 1b) provided refined abundance estimates and enabled direct comparison between cortex and dorsal root ganglia (DRG). In the cortex, consistent with Nanopore data, we observed a similar rank order of TRP transcripts, and found that TRPML, TRPC, and TRPM transcripts collectively account for approximately two-thirds of total TRP expression (Fig. 1C; Supplementary Fig. 3b; Supplementary Table 3). TRPVs were detected at intermediate levels, and TRPP and TRPA were near background under baseline conditions (Fig. 1C; Supplementary Fig. 2b). The cortical TPM estimates included *Mcoln1* (∼65), *Trpc1* (∼20), *Trpm2* (∼20), *Trpm3* (∼21), *Trpm7* (∼13) and *Trpv2* (∼13). In the cortex, the canonical nociceptor channels *Trpa1* (∼0.5) and *Trpv1* (∼0.2) remained near background levels, indicating minimal basal transcription. The read-count-based views from the same libraries were concordant with these abundance estimates (Supplementary Fig. 2b), indicating that the observed per-gene shifts are not attributable solely to length or library-size normalisation. Absolute TPM values varied modestly between the two sequencing platforms, as expected from differences in library preparation and normalisation; therefore, subsequent analyses emphasise rank order and relative enrichment rather than absolute TPM equivalence.

In contrast to the cortex, DRG exhibited a nociceptor-associated TRP profile (Fig. 1D; Supplementary Fig. 2c). In the pooled DRG library, *Trpa1* (∼26) and *Trpv1* (∼37) were among the most abundant transcripts, alongside *Trpc3* (∼47), *Mcoln1* (∼37) and *Trpv2* (∼34), consistent with their established roles in peripheral sensory neurons^15,20,36^. Next, we compared TRP channel gene expression between cortex and DRG, highlighting strong per-gene log_10_(TPM + 1) shifts (Supplementary Figs. 2d,3d). DRG sequencing served as a qualitative positive-control tissue to benchmark detection sensitivity rather than to support statistical differential-expression inference. We observed that *Trpv1* was ∼190-fold higher in DRG than cortex (37.5 vs 0.2), and *Trpa1* was ∼50-fold higher (26.4 vs 0.5); however, these fold differences should be interpreted cautiously because cortical values lie near the detection floor (Supplementary Fig. 3d). On the other hand, *Trpc1* and *Trpm3* were ∼2-fold higher in cortex than DRG (20.2 vs 9.8 TPM and 21.4 vs 9.8, respectively). Collectively, the mouse cortex expresses TRP transcripts from multiple subfamilies but is dominated by TRPML/TRPC/TRPM. In contrast, DRG shows a broader, sensory-enriched repertoire with *Trpa1*/*Trpv1* among the most abundant transcripts.

### TaqMan qPCR validates cortical TRP transcript rankings and cross-platform concordance

To independently validate RNA-seq-derived TRP transcript rankings, we next performed TaqMan qPCR across all 28 mammalian TRP genes in adult mouse cortex and compared these measurements with Illumina and Nanopore RNA-seq. Mean cycle-threshold (Ct; Supplementary Table 4) values were used for descriptive ranking and classified a priori as lower Ct (<27), intermediate Ct (27-32), high Ct/near detection limit (32-40), or not amplified (≥40). The reference gene *Gapdh* amplified robustly (Ct = 17.26), confirming assay performance. Lower Ct values (indicative of higher transcript abundance) were observed predominantly for TRPC and TRPM family members, including *Trpc1, Trpc3–5, Trpm3* and *Trpm7*, as well as a subset of other TRP channels including *Mcoln1*, *Pkd2 (Trpp2)* and *Trpv2* (Fig. 2A). A broader set of TRP genes exhibited intermediate Ct values, whereas several including *Trpa1, Trpm1, Trpm8, Mcoln2, Pkd2l1* and *Trpv1* were detected at high Ct values near the amplification limit; *Mcoln3* and *Trpv5* were not amplified.

**Fig. 2:**
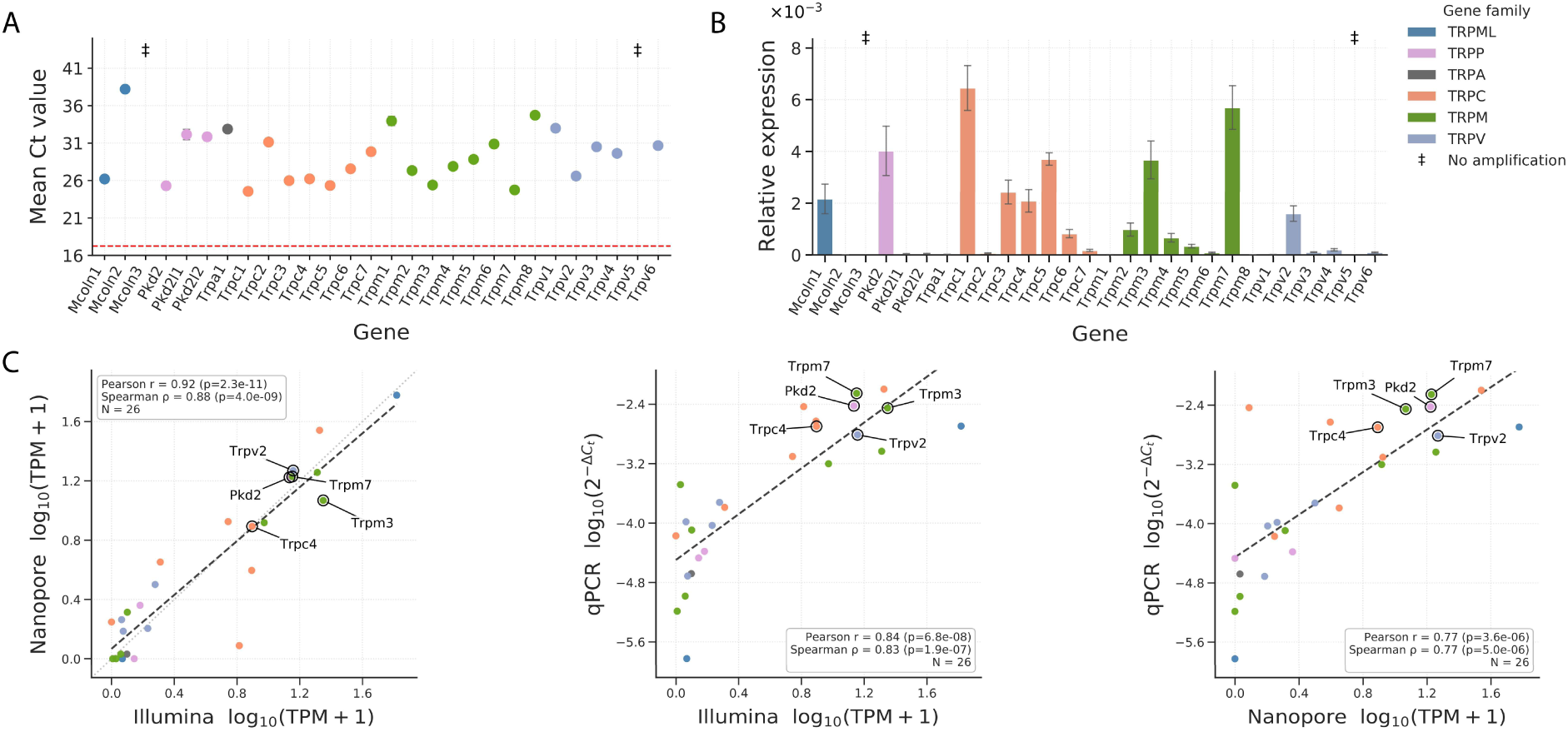
qPCR validation of cortical TRP transcript abundance and cross-platform concordance with Illumina and Nanopore RNA-seq. (**A**) Mean Ct values for TRP genes measured by TaqMan qPCR in adult mouse cortex. Colours denote TRP subfamilies. Red dashed line indicates *Gapdh* Ct. (± SEM; *n* = 3 biological replicates), “‡” denotes no amplification (≥40 cycles; *Mcoln3*, *Trpv5*). (**B**) Relative expression (2^⁻ΔCt^; mean ± SEM, n = 3 biological replicates) after normalisation to *Gapdh* (linear y-axis; external multiplier ×10⁻³). (**C**) Cross-platform concordance among Illumina RNA-seq, Nanopore direct RNA-seq, and *Gapdh*-normalised qPCR measurements for 26 TRP genes. Scatter plots show pairwise comparisons between platforms (log₁₀[TPM + 1] or log₁₀[2⁻^ΔCt^]), with linear regression fits indicated by black dashed lines and Pearson/Spearman correlation coefficients shown within each panel. Selected TRP genes of interest (*Trpv2*, *Trpm3*, *Trpm7*, *Trpc4*, and *Pkd2*) are labelled to aid cross-panel comparison. *Mcoln3* and *Trpv5* were excluded from correlation analyses due to a lack of detectable amplification in qPCR (Ct ≥ 40). Values represent mean expression across three biological replicates (*n = 3*). The identity line (grey dotted) is shown only for the Illumina–Nanopore comparison because both axes use the same units.

Consistent with the high analytical sensitivity of targeted TaqMan qPCR assays, four additional TRP genes (*Mcoln2*, *Pkd2l1*, *Trpm5* and *Trpm8;* Fig. 2A) were detected, which were previously measured below the conservative detection thresholds in bulk RNA-seq (Figs. 1B,C). However, these transcripts were detected only at high Ct values near the assay floor, supporting trace-level expression rather than robust basal transcription. Conversion of ΔCt values (*Gapdh* normalised) to relative expression (2^⁻ΔCt^) closely recapitulated the cortical RNA-seq-derived rank order (Fig. 2B), with TRPC/TRPM transcripts dominating, *Pkd2* and *Trpv2* at intermediate levels, and *Trpa1/Trpv1* at the detection floor.

Cross-platform comparisons demonstrated strong concordance among Nanopore, Illumina and qPCR measurements (Fig. 2C). Illumina and Nanopore RNA-seq abundances (log₁₀[TPM + 1]) were strongly correlated (Pearson r = 0.92; Spearman ρ = 0.88; N = 26, after excluding the two non-amplified genes), with values clustering near the identity line, indicating minimal systematic bias between sequencing platforms. *Gapdh*-normalised qPCR measurements (log₁₀[2⁻ΔCt]) also correlated strongly with Illumina (r = 0.84, ρ = 0.83) and with Nanopore (r = 0.77, ρ = 0.77), despite the narrower dynamic range inherent to qPCR. One gene (*Trpc5*) showed reduced concordance relative to the overall trend, potentially reflecting isoform- or primer-specific detection differences. Together, these analyses demonstrate robust cross-platform agreement and validate the relative ranking of TRP transcripts in adult mouse cortex.

### Membrane-aware proteomics improves recovery of hydrophobic TRPs and defines a restricted cortical TRP proteome

Having established transcript-level TRP expression in adult cortex, we next investigated whether this abundance ranking is reflected at the protein level. TRP channels are multi-pass integral membrane proteins with significant hydrophobic characteristics that complicate conventional mass spectrometric detection^31,33,34^. To overcome this challenge, we systematically evaluated multiple approaches by modifying extraction and purification processes and by incorporating detergent- and urea-based chemistries to enrich membrane fractions (workflow, Fig. 3A; Supplementary Figs. 7a-c; terminology in Supplementary Table 5). These workflows included cytosolic supernatant (SN), crude membrane fractions (Control), urea-solubilised membrane fractions (Pellet), urea-insoluble membrane-enriched fractions (Pelletpellet), filter-aided sample preparation (FASP)-based methods using urea (FASP-8M) or SDS (FASP-SDS), and SDS–PAGE gel-excised samples (Gel). Differences in protein recovery across workflows reflect differences in the solubilisation of hydrophobic and tightly membrane-associated proteins, with urea- and detergent-based approaches generally favouring recovery of integral membrane proteins relative to crude membrane preparations. Proteins were considered detected based on statistically controlled identification rather than raw intensity alone; precursor- and protein-group–level false discovery rates were controlled at 1%, and proteins were required to be identified in at least 2 of 3 biological replicates per workflow (Supplementary Tables 6-7). All samples were analysed using identical LC gradients, data-independent acquisition (DIA) parameters, and data-processing settings to ensure that differences in detection reflect extraction chemistry rather than variation in mass spectrometric depth. Across cortical replicates, multiple unbiased analyses, including clustering of missing-value patterns (undetected proteins), protein identification counts, and PCA, consistently showed high reproducibility within each extraction protocol and clear separation of proteomic profiles across extraction methods (Figs. 3B-D). Membrane-focused workflows, including pellet-derived and FASP-based approaches, reduced missingness and increased proteome coverage relative to supernatant fractions (Figs. 3B-D; Supplementary Figs. 8a,b and 9b). In addition, analysis of global abundance patterns (Supplementary Fig. 9) together with Gene Ontology cellular-component annotations (Supplementary Fig. 10a,b; Supplementary Table 8) indicated that membrane-focused extraction protocols preferentially captured proteins annotated to membrane compartments, with distinct abundance patterns across extraction chemistries consistent with differential protein abundance analyses (Supplementary Tables 9-10).

**Fig. 3:**
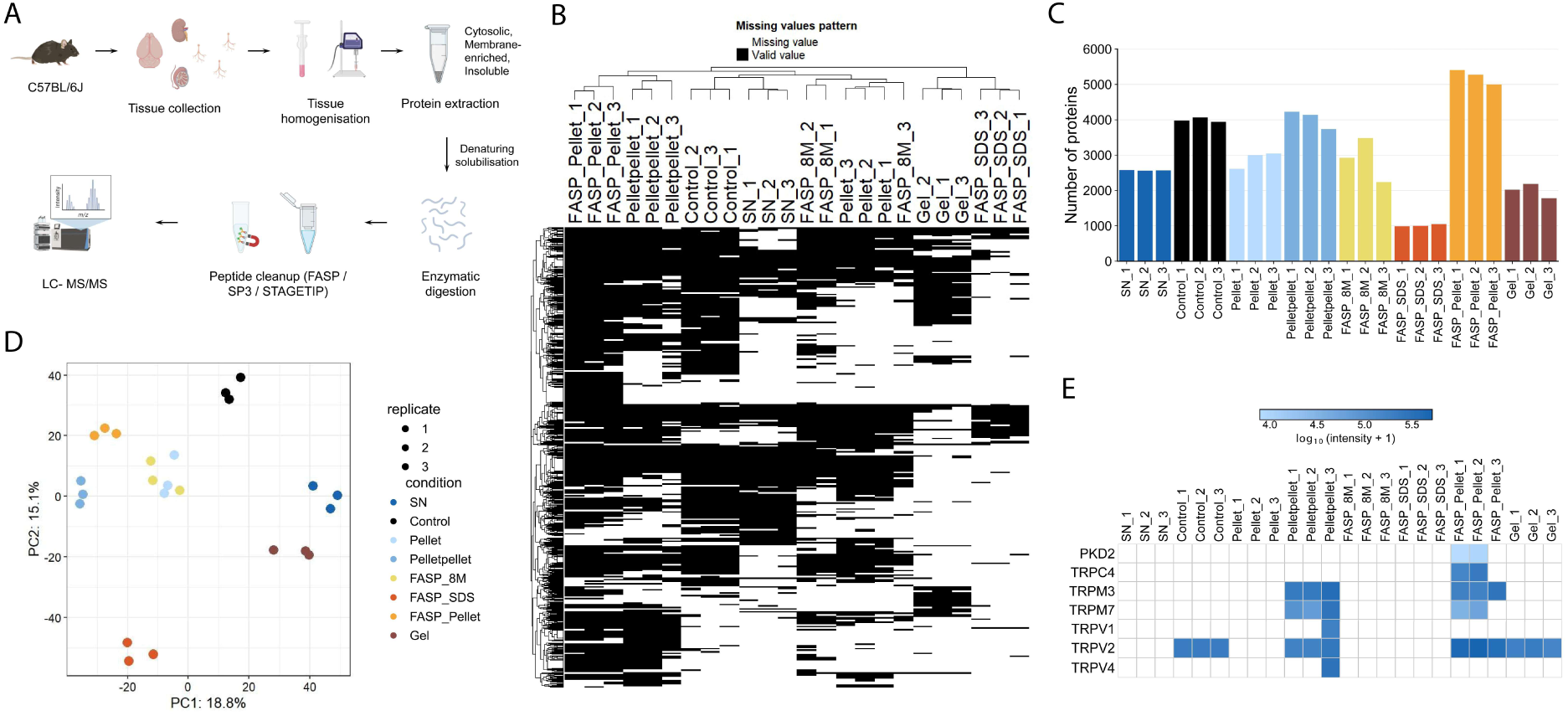
Membrane-aware proteomics enhances recovery of hydrophobic TRP channels. (**A**) Proteomics workflow. Tissues from adult mice were homogenised and fractionated into cytosolic, membrane-enriched, and insoluble fractions. Proteins were solubilised using detergent- or urea-based chemistries, enzymatically digested, cleaned up using FASP, SP3, or StageTip workflows, and analysed by LC–MS/MS. Sample labels used throughout (SN, Control, Pellet and Pelletpellet) correspond to cytosolic, crude membrane, urea-soluble membrane, and urea-insoluble membrane fractions, respectively (see Supplementary Fig. 7; terminology in Supplementary Table 5). (**B**) Missing-value map with unsupervised hierarchical clustering (black = observed; white = missing). Samples cluster by protocol, indicating high within-protocol consistency and distinct missingness profiles across procedures. (**C**) Proteins identified per sample; membrane-enriching inputs (FASP-Pellet, Pelletpellet) yield more identifications than supernatant (SN) or gel-only preparations; replicate variability is low (*n* = 3 per condition). (**D**) PCA of the top 500 variable proteins. Replicates cluster tightly by protocol and separate along PC1/PC2, indicating reproducible and protocol-specific proteome signatures. (**E**) TRP-focused log₁₀-transformed protein intensities [log₁₀(intensity + 1)] heatmap showing evidence for TRP channels (reproducible detection criteria were met for TRPV2, TRPC4, TRPM3, TRPM7 and TRPP2) across protocols. FASP-based and pellet-derived inputs gave the strongest TRP signals. QC: 1% precursor- and protein-group–level detection FDR (DIA-NN) with downstream DEP filtering and imputation.

In line with these observations, a TRP-focused heatmap showed that membrane-enriching workflows recovered more TRP channel peptides and yielded higher TRP protein-group identifications compared to supernatant-only preparations (Fig. 3E; Supplementary Fig. 10c). When analysis was limited to cortical samples (excluding non-cortical tissues and positive controls), and protein-groups were required to pass replicate-level filtering, only a small subset of TRP family members showed consistent protein-level detection. In adult cortical tissue, specifically TRPV2 and the non-canonical TRPM members TRPM3 and TRPM7 were reliably detected at the protein level, predominantly in membrane-enriched workflows. In contrast, the supernatant fractions yielded no reproducible TRP protein-group identifications (Supplementary Figs. 10c,d). TRPC4 and TRPP2 (PKD2) were detected in the cortex but showed more protocol-dependence. TRPV4 was identified at the protein-group level under 1% FDR control but showed low integrated fragment-ion intensity and was observed in only a single biological replicate; therefore, it did not meet replicate-level filtering criteria. By contrast, TRPA1 and TRPV1 did not yield reproducible protein-group–level detection in bulk cortex; only sporadic low-intensity TRPV1 protein-group signals were observed, and these likewise lacked consistent replicate-level support.

To benchmark TRP detection sensitivity beyond the cortex, we analysed non-cortical tissues (DRG, kidney, and testes) together with positive controls (gel-excised SDS–PAGE bands and *Xenopus* oocytes overexpressing rat protein standards). These datasets showed comparable workflow-dependent patterns, including lower missingness in membrane-enriching preparations and distinct sample clustering (Supplementary Figs. 8a,b and 9b), supporting the generalisability of our approach across tissues. As expected, the canonical sensory TRPV1 was readily detected in DRG, and in positive-control preparations (Xenoblot and gel-excise standards; Supplementary Figs. 8c and 12d). TRPV2 was weakly detected in DRG; TRPV4 was observed in kidney and testes pellet preparations, whereas TRPM7 showed broader recovery across DRG and testes pellet samples. A tissue-restricted signal for TRPML3 (MCOLN3) was evident in testes. PKD2 (TRPP2) was detected across DRG, kidney, and testes.

Together, these data demonstrate that our membrane-aware workflows materially improve the detection sensitivity of TRP proteins across conditions. However, we cannot rule out the possibility that some TRP proteins fell below our detection cut-off, not because they are not expressed, but because their peptides are less efficiently recovered or detected by mass spectrometry. Thus, the absence of reproducible detection defines the practical detection limits rather than the definitive absence of protein.

Although TRPV1 and TRPA1 protein expression have been reported in DRG, testes and other tissues in prior studies^37–39^, our analyses did not yield reproducible protein-group–level detection of TRPA1 in any tissue or workflow examined. TRPV1 was reproducibly detected in DRG and in positive-control preparations, but not in cortex, kidney, or testes under the workflows applied here. Such discrepancies may reflect differences in tissue preparation, sensitivity thresholds, antibody-based detection approaches, or biological context. To address these uncertainties directly in the cortex, we next employed complementary cell-type–resolved and immunocapture-based strategies to define stringent abundance bounds for TRPA1 and TRPV1.

### TRPA1 and TRPV1 expression in mouse cortex: Cell-type-specific qPCR, immunoblots, and IP–MS place stringent bounds on abundance

We next asked whether the sensory transducers TRPA1 and TRPV1, expressed in the peripheral ganglia^16,17^, maintain any molecular footprint in the mouse cortex. Given their controversial detection in brain tissue, we combined cell-type-resolved qPCR, fractionated immunoblotting, immunocapture and peptide-level mass spectrometry to identify practical bounds on cortical abundance. Adult cortex was dissociated, and fluorescence-activated cell sorting (FACS) was used to isolate NeuN⁺ neurons, ACSA2⁺ astrocytes, NeuN⁺/ACSA2⁺ double-positive events, and double-negative populations (Supplementary Figs. 11a,b). RNA yields were modest but consistent (∼100 ng total RNA per population), and housekeeping gene expression (*Gapdh* Ct ≈ 23 across fractions; higher than the bulk-cortex *Gapdh* Ct in Fig. 2A; likely reflecting lower input RNA; Ct values in Supplementary Table 4) was consistent, supporting reliable quantification. In bulk cortex, *Trpv1* and *Trpa1* amplified only at high Ct values (≈ 33), indicating near-threshold abundance (Fig. 4A). In sorted populations, *Trpv1* amplification was restricted to NeuN⁺ neurons at a high cycle threshold (Ct ≈ 37) and was not detected in ACSA2⁺ astrocytes, whereas *Trpa1* was only sporadically detected near threshold levels (Ct ≈ 38) in NeuN⁺/ACSA2⁺ double-positive events (Fig. 4B). In contrast, *Mcoln1, Trpc6*, *Trpm2*, *Trpm3* and *Trpv2*, which were used as cortical reference TRPs, amplified ubiquitously across all populations at lower Ct values (Ct 30-35; Supplementary Fig. 11c), confirming the TaqMan assay’s dynamic range and underscoring the scarcity of *Trpa1* and *Trpv1* transcripts.

**Fig. 4:**
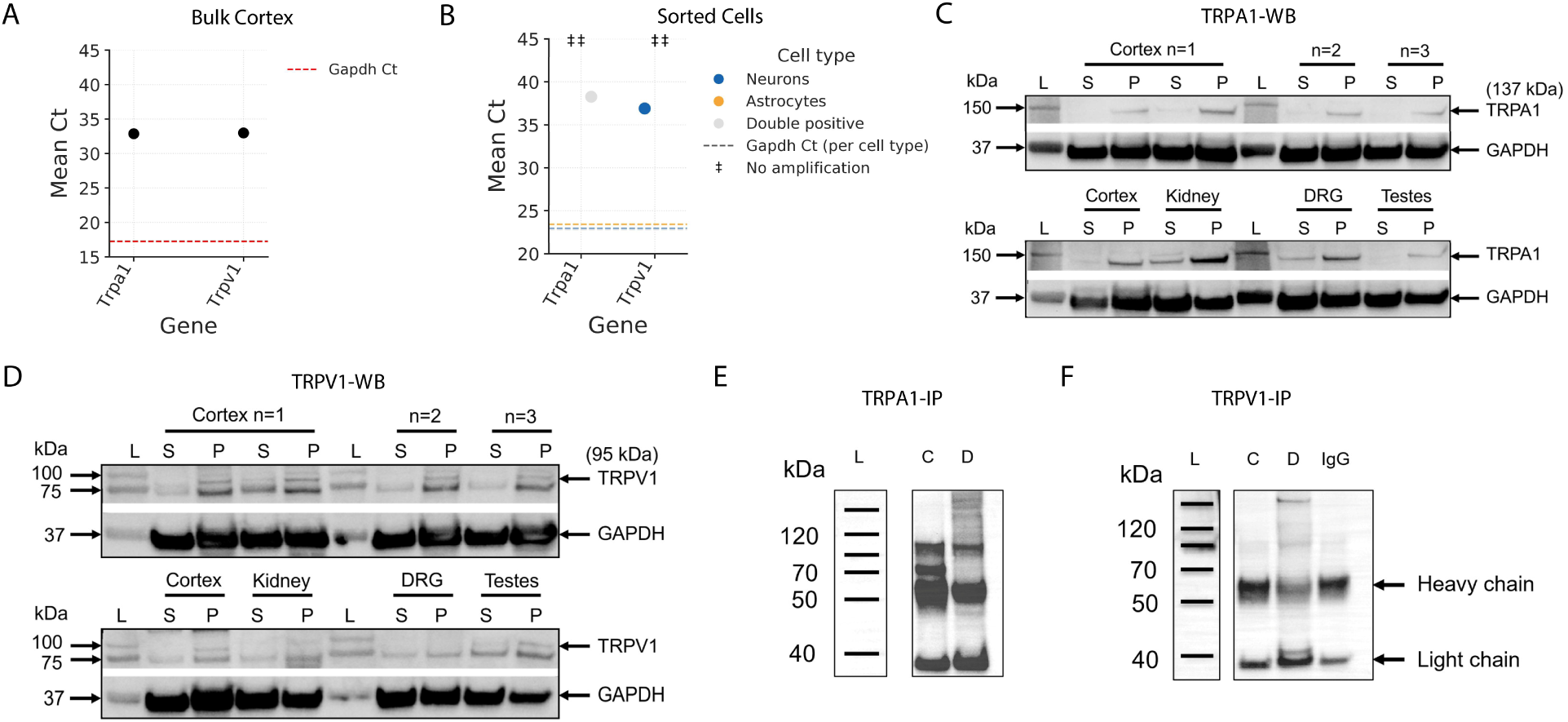
TRPA1 and TRPV1 exhibit near-threshold transcript and protein-level evidence in adult mouse cortex. (**A**) Bulk cortex TaqMan qPCR showing mean Ct values for *Trpa1* and *Trpv1* in adult mouse cortex. Both transcripts are detected only at high Ct values. Red dashed line indicates *Gapdh* Ct. (**B**) TaqMan qPCR of FACS-sorted cortical cell populations showing mean Ct values by fraction: neurons (blue), astrocytes (orange), and NeuN⁺/ACSA2⁺ double-positive cells (grey). Dashed lines indicate fraction-specific *Gapdh* Ct. “‡” indicate no detectable amplification reactions. *Trpv1* is neuron-only at high Ct; *Trpa1* is rare/borderline. (**C, D**) Subcellular fractionation and immunoblotting for TRPA1 and TRPV1, respectively. Cortex lysates (n = 3) were separated into supernatant (S) and membrane pellet (P) fractions; Ladder (L). Faint immunoreactive bands near the expected molecular weight (∼130–140 and ∼95 kDa, respectively) are preferentially enriched in pellet fractions. GAPDH (∼37 kDa) was used as a loading control. Kidney, DRG, and testes processed in parallel serve as positive control tissues. Note: Lysates and controls were processed in the same experiment and run on the same gels/blots. (**E, F**) Immunoprecipitation (IP) from cortex (C) and DRG (D) lysates followed by SDS–PAGE/Western blotting and LC–MS/MS. As shown in Supplementary Fig. 12f, no TRPA1 peptides were detected, and only extremely low-intensity TRPV1 protein-group signals were observed without replicate consistency.

At the protein level, subcellular fractionation of cortex lysates revealed faint immunoreactive bands in the approximate molecular-weight range of TRPA1 (∼130–140 kDa) and TRPV1 (∼95 kDa), preferentially enriched in membrane pellet fractions relative to cytosolic supernatants across biological replicates (Figs. 4C,D), consistent with membrane association characteristic of canonical TRPs. Parallel fractionation of kidney, DRG, and testes, positive control tissues for both channels, showed a similar pellet-enriched immunoreactivity (Figs. 4C,D; Supplementary Figs. 12a-c). GAPDH (∼37 kDa) was uniformly distributed and used as a loading control (Fig. 4D; Supplementary Figs. 12b,c). However, given the low apparent abundance of TRPA1/TRPV1 in cortex (Figs. 4C,D; Supplementary Figs. 13a-d) and potential susceptibility of antibody-based detection to cross-reactivity at low signal levels, these immunoblot data alone could not establish peptide-level specificity. Two independent antibodies targeting TRPA1 produced broadly distributed immunoreactive signals (Supplementary Fig. 13a, c) in cortical sections, suggesting variability in epitope recognition and potential off-target binding under low-abundance conditions. Similarly, TRPV1 antibodies yielded detectable signal in cortical sections (Supplementary Fig. 13b,d), despite the absence of reporter signal in cortex in TRPV1-Cre-eYFP mice (Supplementary Fig. 14), where robust expression was confirmed in peripheral positive-control tissues (DRG, testis, reporter-positive tissues); accordingly, immunofluorescence alone was interpreted cautiously and was not used as standalone evidence of endogenous cortical protein expression.

We therefore sought peptide-level confirmation using immunoprecipitation (IP) from cortical lysates followed by LC–MS/MS (Fig. 4E). Consistent with previous cortex-restricted bulk peptide analyses (Fig. 3E), TRPA1 did not yield protein-group–level detection from cortical IPs under stringent 1% precursor- and protein-group–level FDR control (Fig. 4E). For TRPV1, LC–MS/MS re-analysis incorporating Xenoblot-derived TRPV1 peptide reference (Supplementary Figs. 12d-f; Fig. 4F; Supplementary Table 11) together with DRG positive controls yielded extremely low-intensity protein-group evidence for TRPV1 in a limited subset of cortex IP samples, corresponding to the same peptide sequence without replicate-level consistency. This pattern is consistent with stochastic recovery at or near the detection limit rather than robust cortical TRPV1 expression. Although a recombinant spike-in titration was not performed to define an absolute detection threshold, the absence of reproducible protein-group–level detection across preparations indicates that cortical TRPV1 abundance lies at or below the practical sensitivity of standard LC–MS/MS workflows.

Together, these cell-type transcriptomic and peptide-level data define a stringent molecular boundary for cortical TRPA1 and TRPV1: transcripts are low, and protein abundance remains near or below the practical detection limits of bulk proteomics, even after membrane enrichment and immunocapture, in healthy adult cortex under baseline conditions. In contrast, multiple non-sensory TRP channels were reproducibly detected at the protein level, concordant with transcriptomic prevalence and validating the sensitivity gains achieved through membrane-aware proteomics. Collectively, these multi-layered findings provide a framework for reconciling reported discrepancies between antibody- and transcript-based studies by distinguishing robustly expressed cortical TRPs from canonical sensory channels whose cortical presence, if any, appears exceedingly limited under physiological conditions.

## Discussion

### TRPC and TRPM family members dominate cortical TRP landscape

TRP channels anchor canonical sensory transduction in the periphery, yet their contribution to cortical signalling has remained uncertain. Their molecular footprint in the adult cortex is comparatively sparse and technically difficult to resolve. Here, we addressed two persistent limitations: i) insufficient transcript resolution for low-abundance multi-exon genes and ii) poor proteomic recovery of hydrophobic multi-pass membrane-embedded proteins, often confounded by antibody cross-reactivity at low abundance. By combining short- and long-read RNA-seq, highly sensitive TaqMan assays and membrane-aware LC–MS/MS, we establish a quantitative baseline abundance ranking of TRP channel expression in the adult mouse cortex and define practical protein-group detection boundaries under the applied workflows. Our multi-platform strategy mitigates the biases inherent to any single method.

Across our Nanopore, Illumina, and TaqMan datasets, cortical TRP expression was dominated by TRPML, TRPC, and TRPM family members, with strong cross-platform concordance. *Mcoln1 (Trpml1), Trpc1, Trpm2/3/4/7* consistently ranked among the most abundant cortical TRPs, while *Trpv2* was also reproducibly detected at relatively high abundance. By contrast, the archetypal nociceptor channels *Trpa1* and *Trpv1* transcripts were detected near the limit of detection under physiological conditions (Fig. 1B, C; Fig. 2A, B). DRG exhibits the inverse pattern, with robust *Trpa1* and *Trpv1* expression and relative attenuation of cortical-enriched TRPs (Fig. 1D; Supplementary Fig. 2d), emphasising a tissue-specific, nociceptor-enriched TRP repertoire in peripheral ganglia. This divergence suggests that cortical TRP biology is organised around signalling pathways distinct from peripheral nociceptive transduction, and that “sensory TRP” assumptions do not generalise across tissues. Instead, the enrichment of TRPC and TRPM channels points to a cortical TRP landscape centered on Ca²⁺ homeostasis, metabolic sensing, and excitability within intrinsic circuits, consistent with established roles for *Trpc4*, *Trpm3*, *Trpm4*, and *Trpm7* in dendritic Ca²⁺ entry, astrocyte–neuron signalling and activity-dependent membrane dynamics^40–42^.

Broadly, the cortical transcript-level hierarchy observed here aligns with earlier RT-PCR-based profiling on rodent whole brain, reporting enrichment of *Trpm3* and *Trpm7*^39,43^, and extends these findings with isoform-resolved and protein-level validation. By reconstructing full-length transcripts rather than aggregating gene-level counts, we identify low-abundance, cortex-enriched TRP variants and refine subfamily-specific resolution (Fig. 1B; Supplementary Fig. 4) within large-scale transcriptomic surveys^25,44^. Independent single-cell and single-nucleus atlases similarly localise TRPC and TRPM members (*Trpc1*, *Trpc4*, *Trpm3*, *Trpm4*, *Trpm7*) to defined neuronal and glial subclasses, while *Trpv2* associates with neuronal, vascular and microglial populations^27,45–47^. Human datasets reveal regional and cell-type gradients in orthologous TRP genes that differ from those in the mouse cortex^24^, suggesting species-specific TRP hierarchies and potentially divergent CNS pharmacological responses. Together with prior reports of strain-dependent peripheral TRP variation^39,48^, the present data define a *C57BL/6J*-specific cortical baseline that should not be assumed to generalise across species or genetic backgrounds.

### Membrane-aware proteomics defines practical detection boundaries

A key technical advance of this study is the demonstration that protein extraction chemistry critically determines the detectability of hydrophobic channels by LC–MS/MS. Using workflows optimised for membrane proteins (detergent, urea-tolerant and membrane-enriching workflows), we reproducibly detected peptides from several TRP channels in adult cortex, including TRPV2, TRPC4, TRPM3, TRPM7, and TRPP2 (Fig. 3E; Supplementary Fig. 8c), confirming their steady-state protein presence in concordance with transcript-level rank order. Reproducible protein-group–level quantification across biological replicates under stringent FDR control establishes an empirically constrained boundary for steady-state protein detectability under the applied DIA workflow. Within this framework, non-detection or isolated single-peptide matches are interpreted as quantitative constraints rather than evidence of absolute absence.

TRPA1 was not detected at the protein-group level in cortex, DRG, kidney, or testes under identical membrane-aware workflows, despite transcript-level enrichment in DRG (Fig. 3E; Supplementary Fig. 8c). TRPA1 likewise did not yield robust protein-group–level signals from the Xenoblot preparation under denaturing conditions. Although DIA acquisition is designed to sample precursor ions comprehensively, protein detectability remains constrained by peptide recovery, ionisation efficiency, and tissue complexity^49^. Similar analytical challenges have been reported for Ankyrin-rich TRPA1 in membrane-focused DRG proteomics, despite established nociceptor expression^50–52^. Moreover, TRPA1 expression in sensory neurons may not be strictly constitutive under basal conditions and can vary heterogeneously across ganglia, depending on prior cellular state, injury paradigm or nociceptive stimulation^53–55^. TRPV1 yielded only sporadic, low-intensity protein-group signals in the cortex, without consistent replicate-level support. In contrast, it was reproducibly detected in DRG but not across other peripheral tissues. Together, these observations indicate that steady-state TRPA1 and TRPV1 protein abundance in adult cortex is substantially lower than that of TRP family members reproducibly detected across replicates.

Disparities between TRP transcript and protein abundance in the cortex are expected and analytically informative. For example, *Mcoln1* transcripts rank among the most abundant cortical TRPs, yet the protein was not reproducibly recovered. As a lysosomal multi-pass channel, TRPML1 proteomic recovery may be limited by subcellular compartmentalisation, hydrophobic peptide bias and state-dependent lysosomal activation/expression^56^. More broadly, many membrane and synaptic proteins in post-mitotic neurons exhibit slow turnover^28,30,57^, yielding low steady-state peptides despite measurable transcripts. From a technical perspective, label-free proteomics systematically under-represents highly hydrophobic, transmembrane-rich, ultra-low-copy proteins due to poor solubility and tryptic peptide bias^32,34,58,59^, limitations that our membrane-aware workflows substantially mitigate but cannot fully overcome.

### Reconciling conflicting evidence for cortical TRPA1 and TRPV1

Reports of cortical *Trpa1* and *Trpv1* range from “rare” to “functionally relevant,” with inferences strongly dependent on assay and cellular context. Single-cell atlases and bulk transcriptomic resources consistently report very low transcript abundance, with sparse protein detection^25,27,45^. Nonetheless, pharmacological and genetic studies have implicated their roles in cortical excitability and sensory processing^10,60,61^, including astrocyte-mediated regulation of basal Ca²⁺ levels, inhibitory synaptic efficacy, and gliotransmission^62,63^. Discrepancies likely arise from antibody-based detection, which is vulnerable to cross-reactivity, particularly at low abundance^37,64–73^. Transgenic reporters may reflect transient developmental expression that falls below adult immunohistochemical thresholds^37^, radioligand assays lack cellular resolution^69^ or inadvertently cross-detect related proteins^70^ and knockout-controlled electron microscopy, while specific, remains region-limited^74^. Together, these constraints have complicated efforts to determine the physiologically relevant molecular baseline footprint of TRPA1 and TRPV1 in the mammalian cortex.

Our orthogonal approach places stringent quantitative bounds on cortical TRPA1 and TRPV1 abundance. First, bulk RNA-seq and targeted qPCR consistently position both transcripts near the detection floor (Figs. 1B,C and 2A,B, and 4A). Second, FACS-qPCR indicates that if *Trpv1* is present, it is rare and neuron-skewed, while *Trpa1* appears sporadic and near-threshold in sorted fractions (Fig. 4B), establishing upper bounds on cellular prevalence. While sorting focused on neurons and astrocytes, any rare non-neuronal expression would likely remain diluted below bulk detection thresholds. Third and most decisively, peptide-level proteomics fail to detect TRPA1 and yields only isolated, non-reproducible TRPV1 peptide matches (Figs. 3E and 4E,F). Taken together, the most parsimonious inference is that endogenous cortical TRPA1/TRPV1 are extremely rare at baseline in healthy adult cortex.

Prior reports suggesting context-dependent expression in restricted loci or after defined perturbations raise the possibility of these channels becoming more detectable under such settings^66,68,75^; however, in the absence of strong and reproducible protein-level evidence, such interpretations remain provisional. Candidate loci include the ventricular ependyma^76^, arterial/capillary endothelium mediating neurovascular coupling^77–79^, and microglial or perivascular compartments. Perturbations such as pain^38,80–82^, seizure^83,84^, inflammation^68,85^, neurodegeneration^86^, or injury^87^ may induce cortical/hippocampal TRPA1/TRPV1 expression, potentially fine-tuning local circuit excitability. This model reconciles prior functional observations with the quantitative constraints imposed by bulk transcriptomics and peptide-level proteomics.

## Conclusion

This cross-platform, membrane-aware framework establishes reproducible quantitative baselines for cortical TRP expression and refines the molecular landscape beyond binary detection claims. TRPML, TRPC and TRPM members dominate the basal cortical repertoire, consistent with roles in Ca^2+^ homeostasis and synaptic modulation. The canonical sensory channels TRPA1 and TRPV1 lie at the practical limits of detection in healthy adult cortex, with near-threshold transcripts and no reproducible protein-group–level detection under standard DIA-LC–MS/MS conditions. These results support a conditional-expression model in which TRPA1/TRPV1 may become detectable primarily under perturbations such as seizures, inflammation or injury, thereby refining the translational window for TRP-targeted therapeutics^11,68,84^.

More broadly, given low abundance and sequence homology within TRP families, pharmacological studies must carefully consider ligand specificity and cross-reactivity, as many commonly used TRP modulators engage multiple TRP and non-TRP targets at physiologically relevant concentrations. Moving forward, knockout-validated monoclonal antibodies, together with orthogonal protein measurements, will be essential to localise rare TRPs suggested by our transcriptomic and proteomic data. Overall, this integrated molecular census of cortical TRPs prioritises specific targets for hypothesis-driven experiments in epilepsy and related disorders and provides a scalable framework for investigating low-abundance ion channels within complex brain tissues.

## Methods

### Animals

All procedures used C57BL/6J adult male mice (5–6 weeks old), group-housed at the Australian Phenomics Facility (The Australian National University) under a 12:12 h reverse light/dark cycle (22 ± 2°C) with food and water *ad libitum*. Mice were selected from available age-matched animals housed under identical conditions. In addition, adult male TRPV1-Cre-eYFP mice of a comparable age were used for immunofluorescence experiments (Supplementary Fig. 14); this reporter line was imported from Professor Brett Graham (The University of Newcastle, Australia) and maintained at the Australian National University. All experimental protocols were approved by the ANU Animal Experimentation and Ethics Committee (A/2023/304) and complied with the NHMRC Australian Code for the Care and Use of Animals for Scientific Purposes and the ACT Animal Welfare Act 1992.

### Brain dissection and RNA extraction

Mice were deeply anaesthetised with isoflurane and transcardially perfused with ice-cold artificial cerebrospinal fluid (aCSF; composition in mM: 125 NaCl, 25 NaHCO_3_, 3 KCl, 1.25 NaH_2_PO_4_, 25 Glucose, 2 mM CaCl_2_, 1 mM MgCl_2_, osmolarity adjusted to 300 –310 mOsm and pH adjusted to 7.2 –7.3 with carbogen bubbling: 95% O₂/5% CO₂) before euthanasia. Brains were rapidly extracted within 2 min, rinsed in ice-cold phosphate-buffered saline (PBS, pH 7.4), and placed on a chilled Petri dish for dissection. Bilateral cortical tissue was dissected under a stereomicroscope (Wild Heerbrugg) on ice. Total RNA was extracted by homogenising cortical tissue in 500 µL TRI Reagent (MilliporeSigma, T9424) and purified using the Direct-zol™ RNA Microprep Kit (Zymo Research), including on-column DNase I treatment (Thermo Scientific, EN0521), according to the manufacturer’s instructions.

Dorsal root ganglia (DRG) were rapidly dissected from adult mice following perfusion under the same conditions described for cortical tissue, as previously described^88^. Ganglia were collected bilaterally from the spinal column, pooled where indicated, and immediately processed or snap-frozen on dry ice for downstream analyses. Kidney and testes samples were collected using standard dissection procedures and processed in parallel with cortical tissue.

### Nanopore direct RNA library preparation and sequencing

Direct RNA libraries were prepared with the Oxford Nanopore Direct RNA Sequencing Kit (SQK-RNA004) following the manufacturer’s protocol, substituting Induro Reverse Transcriptase (NEB, M0681) for the kit RT and adding 1 µL SUPERase·In during adaptor ligation. Two libraries were sequenced on MinION R9.4.1 flow cells and one on a PromethION FLO-PRO004RA flow cell, all operated with MinKNOW v24.06.10. Flow cells were equilibrated to 25 °C, pore health was verified by a flow-cell check, and sequencing proceeded to pore exhaustion; raw signals were recorded in POD5 format.

### Reference genomes

Reference genome assemblies and annotations were obtained from Ensembl release 115 for *Mus musculus* (GRCm39). Long-read genome alignment and isoform discovery (IsoQuant) used the GRCm39 unmasked primary assembly (FASTA) together with the corresponding Ensembl r115 gene annotation (GTF). Long-read transcriptome-based quantification (NanoCount) used the Ensembl r115 cDNA FASTA (*Mus_musculus.GRCm39.cdna.all.fa*) with transcript-to-gene mappings from the matching r115 GTF. Short-read quantification (Salmon) used a decoy-aware index built from the Ensembl r115 cDNA plus primary-assembly decoys. Unless stated otherwise, all gene identifiers/symbols are from Ensembl r115, and all genomic coordinates refer to the GRCm39 primary assembly.

### Basecalling and QC of nanopore data

Basecalling and alignment were performed on compute nodes equipped with Tesla Volta V100-SXM2-32GB GPUs. Raw signals (POD5 files) were basecalled using Dorado v0.7.0 with poly(A) tail estimation enabled (--estimate-poly-a), generating unmapped BAM files; FASTQ files were extracted using samtools v1.22 (samtools fastq -T “*”) for downstream analyses. Run-level sequencing summaries, including read yield and N50, were obtained from final run reports, and read-level quality control was further evaluated using NanoPlot. Across the three biological replicates of nanopore direct RNA sequencing, replicates 1, 2, and 3 generated 862,630, 1.32 million, and 8.38 million reads, respectively, with read N50 values of 1.22 kb, 2.72 kb, and 2.49 kb. Alignment of reads to the *Mus musculus* GRCm39 reference genome using minimap2 produced overall mapping rates of 86.91%, 76.00%, and 79.25% for replicates 1, 2, and 3, respectively.

### Gene-level quantification (NanoCount)

Basecalled Nanopore direct RNA reads from adult mouse cortex were aligned to the *Mus musculus* GRCm39 reference transcriptome (Ensembl release 115; *Mus_musculus.GRCm39.cdna.all.fa*) using minimap2 v2.28 with Oxford Nanopore–optimised parameters (-ax map-ont -p 0.8 -N 50). Resulting alignments were coordinate-sorted and indexed using Samtools. Transcript-level abundance estimation was performed independently for each biological replicate using NanoCount v1.2.1 with default settings. NanoCount excludes low-quality alignments (including unmapped reads, invalid 3′ ends, very short or supplementary alignments) and resolves multi-mapping reads by expectation–maximisation to assign reads across compatible transcripts, yielding per-transcript estimated counts and transcripts-per-million (TPM) values.

For gene-level expression summaries, Ensembl transcript identifiers were mapped to genes using the Ensembl r115 gene annotation (*Mus_musculus.GRCm39.115.gtf*). Transcript TPM values were summed per gene within each replicate, and gene-level TPMs were averaged across the three cortex replicates for downstream analyses and visualisation. Gene-level TPMs and read counts from Nanopore RNA-seq are provided in Supplementary Table 1. Low-abundance signals were interpreted conservatively; TPM values below 0.1 were considered effectively indistinguishable from background and below the limit of reliable detection, whereas values between 0.1 and 1.0 were classified as near-background expression.

### Isoform discovery and quantification (IsoQuant)

Basecalled ONT direct-RNA reads from three biological replicates were aligned to the *Mus musculus* GRCm39 primary assembly using minimap2 v2.28 in spliced mode optimised for ONT direct RNA sequencing (-ax splice -uf -k14 --secondary=no). Alignments were coordinate-sorted and indexed using samtools v1.22. Isoform modelling and structural classification were performed with IsoQuant v3.7.0 (ONT mode) using the GRCm39 genome FASTA and Ensembl release 115 gene annotation (GTF) as a guide, without restricting transcript discovery to annotated models. IsoQuant collapsed long reads into non-redundant transcript models, assigned models to genes, and generated SQANTI-like structural classifications, including full-splice-match (FSM/known), novel-in-catalog (NIC), and novel-not-in-catalog (NNC) categories.

Isoform abundance was quantified per replicate as transcripts per million (TPM), and gene-level TPM values were calculated as the sum of constituent isoforms. Known and novel TRP isoforms identified in the adult mouse cortex are listed in Supplementary Table 2. For downstream analyses, an isoform was considered present if it had non-zero TPM in at least 2 of the 3 biological replicates. This criterion was used to prioritise isoforms reproducibly detected across biological replicates while avoiding arbitrary abundance cut-offs for long-read data. Novel transcript models highlighted in Supplementary Fig. 4 were further inspected in IGV using the processed BAM files to verify splice-junction support by junction-spanning reads with canonical splice motifs, confirm consistent exon connectivity, and exclude rare chimeric alignments.

### Illumina RNA library preparation and sequencing

Total RNA (2 µg per sample) from mouse cortex and DRG was used to generate strand-specific libraries using the NEBNext® Ultra™ II Directional RNA Library Prep Kit (New England Biolabs), following poly(A)+ selection with the NEBNext® Poly(A) mRNA Magnetic Isolation Module (E7490). For DRG, tissue from three mice was pooled per library (n = 1) to maximise RNA yield and sensitivity. Purified mRNA was fragmented and reverse-transcribed, followed by second-strand synthesis, end repair and A-tailing, adaptor ligation, and PCR amplification (12 cycles). Libraries were quantified by qPCR (KAPA SYBR® Fast, KK4601), pooled equimolarly, and sequenced by Azenta Life Sciences on an Illumina NovaSeq 6000 platform to generate 150-bp paired-end reads (PE150). Library quality control included Qubit fluorometric quantification and fragment size assessment using a Bioanalyzer (Agilent 2100/2200).

### Read processing and reference annotations

Primary basecalling was performed on the instrument using Illumina RTA software, and demultiplexing was carried out by the sequencing provider using bcl2fastq v2.20, generating paired-end FASTQ files. All downstream analyses used the *Mus musculus* reference genome GRCm39 and Ensembl release 115 annotations. Sequencing generated 76.1 million reads for cortex-rep1, 67.6 million reads for cortex-rep2, 47.8 million reads for cortex-rep3, and 57.8 million reads for the pooled DRG library, with Q30 values of 85.9%, 86.6%, 85.5%, and 86.5%, respectively.

FASTQ files were quality-filtered and adapter-trimmed using FASTX-Toolkit v0.0.14. Reads were filtered with fastq_quality_filter to retain sequences in which at least 40% of bases had a minimum Phred quality score of 33. Adapter clipping was performed using fastx_clipper with Illumina adapter sequences (Read 1: AGATCGGAAGAGCACACGTCTGAACTCCAGTCAC; Read 2: AGATCGGAAGAGCGTCGTGTAGGGAAAGA), and reads shorter than 35 bp after trimming were discarded. Paired-end read synchronisation was enforced using fastq-pair, and only properly paired reads were retained for downstream analysis.

### Quantification (Salmon)

Transcript abundance was quantified using Salmon v1.10.1 in quasi-mapping mode against a decoy-aware GRCm39 transcriptome index (k-mer = 25). Quantification was performed with the parameters --libType A --validateMappings --seqBias --gcBias. Gene-level estimates were generated during quantification using --geneMap with an Ensembl r115 transcript-to-gene mapping file, which recalculates effective lengths and reports gene-level TPM values.

Transcript- and gene-level estimates were aggregated across biological replicates for downstream analyses. Gene-level TPMs and read counts from Illumina RNA-seq are provided in Supplementary Table 3. TRP channel genes were extracted from gene-level quantifications, and expression distributions (TPM and counts) were summarised and visualised in Python.

### TaqMan qPCR assay

Total RNA (2 µg per reaction; DNase-treated) from mouse cortex was reverse-transcribed using SuperScript™ II Reverse Transcriptase (Thermo Fisher Scientific) in a 20 µL reaction following the manufacturer’s instructions, with random hexamer priming and RNase inhibitor. Briefly, RNA was combined with random hexamers (50–250 ng; 1 µL), dNTP mix (10 mM each; 1 µL), and nuclease-free water to 12 µL, heated to 65 °C for 5 min, and snap-chilled on ice. The following reagents were then added: 5× First Strand Buffer (4 µL), 0.1 M DTT (2 µL), and RNaseOUT™ (40 U/µL; 1 µL). After gentle mixing, the reaction was incubated at 25 °C for 2 min. SuperScript™ II (200 U; 1 µL) was added, mixed, and incubated at 25 °C for 10 min, followed by 42 °C for 50 min. The enzyme was inactivated at 70 °C for 15 min, and cDNA was stored at −20 °C until use.

TaqMan assays were performed on a QuantStudio™ 12K Flex (Applied Biosystems, Thermo Fisher Scientific) using TaqMan™ Fast chemistry. Each 20 µL reaction contained 10 µL TaqMan™ Fast Advanced Master Mix (2×), 1 µL TaqMan™ Gene Expression Assay (20×; assay IDs in Supplementary Table 4), 3 µL cDNA (undiluted), and 6 µL nuclease-free water. Plates were briefly vortexed, centrifuged, and sealed; no-template controls showed no amplification. All samples were run in technical duplicates with three biological replicates. Cycling (fast mode): 50 °C for 2 min (UNG), 95 °C for 20 s, then 50 cycles of 95 °C for 1 s and 60 °C for 20 s with fluorescence acquisition at 60 °C. Ct values for the whole cortex and FACS-sorted populations are provided in Supplementary Table 4. Ct values were called automatically, and relative expression was calculated by the 2^−ΔCt^ method using *Gapdh* as the reference. Technical duplicates were averaged to obtain a single Ct value per biological replicate prior to ΔCt calculation and statistical analysis.

### Cell suspension preparation and Fluorescence-activated cell sorting (FACS)

Adult mouse cortical tissue was dissected following perfusion with ice-cold aCSF to remove erythrocytes. Cortices were isolated on ice, excluding cerebellum, striatum, hippocampus, and piriform cortex, and finely minced (1–2 mm³) under oxygenated conditions. Cortical tissue from adult mice was dissociated into a single-cell suspension using a combination of mechanical and enzymatic methods. For enzymatic dissociation, a papain digestion solution was prepared in a carbogen-equilibrated base buffer containing (final concentrations, in mM): NaCl 92, KCl 2.5, NaH₂PO₄ 1.2, HEPES 20, glucose 25, sodium pyruvate 3, MgSO₄ 2, CaCl₂ 1, and NaHCO₃ 30. The base buffer was adjusted to pH 7.35–7.45 and ∼310 mOsm before the addition of enzymes. Immediately before digestion, the following additives were included: L-cysteine (2 mM), DNase I (50 U/mL; RNase-free), polyvinyl sulfonic acid (PVSA, 0.30 mg/mL; Sigma) and CNQX (100 µM). Papain (24 U/mL; Sigma P4762) was added last to the cysteine-containing digestion buffer and allowed to pre-activate for 5–10 min at 34–35 °C. The minced cortical tissue was then digested in this solution for 15–20 min at 34–35 °C with gentle mixing (200–300 rpm; Eppendorf Thermomixer C), with progress monitored every 5 min. No EDTA was present during DNase exposure. Digestion was terminated by adding an equal volume of pre-chilled PBS (4 °C).

Mechanical trituration was performed with fire-polished Pasteur pipettes of decreasing tip size until a single-cell suspension was obtained. Myelin and debris were removed using a myelin removal kit (Miltenyi Biotec; #130-096-733) according to the manufacturer’s instructions, and samples were filtered through 500 µm and 70 µm strainers. Cells were pelleted (300–400 × g, 5 min, 4 °C) and resuspended in PBS supplemented with 50 U/mL DNase I and 0.02–0.04% BSA. For live cell staining, cells were incubated with fluorophore-conjugated antibodies (1 in 200 dilution of Anti-NeuN-Alexa488 and Anti-ACSA2-APC; BD Biosciences) for 20–30 min at 4 °C in the dark, washed once in FACS buffer (PBS, 0.02–0.04% BSA and 50 U/mL DNase I), and resuspended gently. Dead cells were excluded by adding DAPI (1.43 µM).

Accutase (Thermo 00-4555-56) was added before sorting to reduce cell clumping and clogging of the FACS nozzle. Cells were sorted on a BD FACSAria or BD FACSFusion using a 70 µm nozzle at 45 psi. Sorting gates were defined on forward/side scatter (FSC/SSC) and fluorophore intensity, with compensation controls applied to correct any spectral overlap. Sorting was performed in purity mode. Sorted cells were collected directly into 1.5–2 mL tubes preloaded with 500 µL TRIzol LS (3:1 ratio sample: TRIzol). Post-sort purity was assessed by re-analysing a small aliquot of sorted cells in PBS. RNA was extracted using the Zymo Direct-zol RNA Microprep kit with on-column DNase digestion, following the manufacturer’s instructions. The RNA from sorted single- and double-positive populations was quantified using a Nanodrop spectrophotometer and reverse-transcribed for downstream TaqMan PCR.

### Cytosolic and membrane protein extraction

Cortical tissue was dissected from adult mice aged 5–6 weeks as described above. Protein extraction was performed using the Thermo Scientific Mem-PER™ Plus Membrane Protein Extraction Kit (#89842) according to the manufacturer’s protocol. The extracted proteins were used for both mass spectrometry and western blotting. Briefly, 20-40 mg of freshly dissected cortex was placed in a 5 mL low-protein bind microcentrifuge tube and washed with 4 mL of cell wash solution. After brief vortexing, the wash was discarded. Homogenisation was performed by adding 500 µL of permeabilisation buffer supplemented with Halt™ protease and phosphatase inhibitor cocktail (Thermo #78442; 1X final concentration). The cortex samples were minced and homogenised with 6-10 strokes at low speed to minimise foaming, with Bel-Art® disposable pestles (Merck BAF199230001) directly in the tube. An additional 500 µL of permeabilisation buffer with protease and phosphatase inhibitors was added, and samples were incubated for 10 min at 4 °C with constant mixing. Cytosolic proteins were separated by centrifugation at 16,000 × g for 15 min at 4 °C, and the supernatant was collected and kept on ice for processing or stored at −80 °C for later experiments. The remaining pellet was resuspended in 1 mL of solubilisation buffer (containing protease and phosphatase inhibitors), homogenised by pipetting, and incubated for 30 min at 4 °C while constantly mixing. A second centrifugation at 16,000 × g for 15 min at 4 °C yielded the membrane fraction in the supernatant, which was either processed immediately or stored in aliquots at −80 °C to avoid freeze-thaw cycles.

### Preparation of transmembrane protein fractions

Urea-soluble membrane proteins: Membrane pellets were precipitated by adding 4 volumes (∼400 µL) of ice-cold acetone and incubated overnight at −20 °C. Pellets were collected by centrifugation at 15,000 × g for 15 min at 4 °C and resuspended in 200 µL of 8 M urea in 1× TBS to enrich integral membrane proteins^89^. After 10 min of sonication on ice, samples were centrifuged again at 15,000 × g for 15 min. The supernatant (urea-soluble proteins) was used for further mass spectrometry analysis. Protein concentration was adjusted to 10 µg in 100 µL (20 mM ABC), followed by reduction with 25 µL of 25 mM TCEP (20 min, 60 °C, 900 rpm) and alkylation with 25 µL of 120 mM IAA (10 min, room temperature, dark). SP3 cleanup and digestion were then performed as follows. An aliquot of urea-soluble proteins was also processed by filter-aided sample preparation (FASP) using published methods^90^ with minor modifications. Protein concentrations were adjusted to 15 µg in 100 µL, and the samples were concentrated using a 50 kDa MWCO filter at 15,000 × g for 15 min. Concentrated proteins >50 kDa were washed with 8 M urea in 1× TBS, centrifuged at 15,000 × g for 15 min, resuspended in 100 µL of 8 M urea in 1× TBS, reduced with 25 µL of 25 mM TCEP (20 min, 60 °C, 900 rpm), and alkylated with 25 µL of 120 mM IAA (10 min, room temperature, dark). Reduced and alkylated proteins were concentrated at 15,000 × g for 15 min, resuspended in 100 µL of trypsin solution (1 µg per 60 µg input protein), and incubated overnight at 37 °C with agitation. Digested peptides were collected by centrifugation at 15,000 × g for 10 min, followed by two washes of the filter unit (50 µL 0.5 M NaCl; centrifugation at 15,000 × g for 10 min), and the combined peptide flow-through was acidified with trifluoroacetic acid (TFA) to a final concentration of 1% prior to peptide desalting as described below.

Urea-insoluble membrane proteins: An aliquot of urea-insoluble protein pellet was resolubilised in 1× TBS containing 4% SDS, and protein concentrations were adjusted to 15 µg in 100 µL. These were processed using the FASP method described above. A subset of samples was treated with PNGase F (A39245; Gibco; digested according to manufacturer’s instructions) following initial filter-aided removal of SDS/urea buffer in an effort to reduce potential steric hindrance. These samples were otherwise processed using the FASP described above. The remaining urea-insoluble pellet was directly processed by adding 25 µL of 25 mM TCEP and reducing at 60 °C for 20 min with agitation. Alkylation was performed with 25 µL of 120 mM IAA for 10 min in the dark. Proteins were digested by adding 100 µL of trypsin solution (1 µL per 40 µg protein), followed by 30-second sonication and overnight incubation at 37 °C with agitation. Digested peptides were centrifuged at 15,000 × g for 10 min, and the supernatant was acidified with TFA to a final concentration of 1% before peptide desalting as described below.

### Protein sample preparation using SP3 Protocol

Protein lysates for cytosolic, membrane, urea-soluble and insoluble fractions were prepared for mass spectrometry analysis using the single-pot, solid-phase-enhanced sample preparation (SP3) method as described by Hughes, et al. ^91^. Briefly, 10 μg of protein was diluted in 100 μL of 20 mM ammonium bicarbonate (ABC) in a 1.5 mL low-protein-bind microcentrifuge tube. Reduction was performed by adding 25 μL of 25 mM tris (2-carboxyethyl) phosphine in ABC, followed by incubation at 60 °C for 20 min with agitation at 900 rpm. Alkylation was achieved by adding 25 μL of 120 mM iodoacetamide in ABC, and the mixture was incubated at room temperature in the dark for 10 min with agitation.

For protein cleanup, 10 μL of a 1:1 mixture of carboxylate-modified paramagnetic speed beads (250 μg each; Cytiva 65152105050250 and 45152105050250) was added to the sample, which was then mixed thoroughly. To enhance protein binding to the beads, 170 μL of absolute ethanol was added, followed by incubation in a Thermomixer C at 24 °C for 5 min with shaking at 900 rpm. The beads were immobilised using a magnetic rack, and the supernatant was discarded. The beads were washed twice with 180 μL of 80% ethanol to remove contaminants.

Proteins bound to the beads were digested by adding 100 μL of trypsin solution (1:40 enzyme-to-substrate ratio). The bead suspension was sonicated for 30 s in a water bath and incubated overnight in Thermomixer® C at 37 °C with shaking at 900 rpm. After digestion, the sample was centrifuged at 15,000 × g for 2 min, magnetically cleared, and the supernatant containing peptides was collected.

### Peptide desalting and Mass spectrometry

All peptide samples were acidified to 1% TFA and were desalted using StageTips. Each StageTip was prepared by inserting three 5-μg SDB-RPS membrane plugs (Sigma #66886-U; 0.8–1.2 mm punch size) into a 200 μL pipette tip. The tips were first conditioned with 100 μL of 100% acetonitrile (ACN) and centrifuged at 1000 × g for 1 minute. This was followed by equilibration with 100 μL of 30% methanol containing 1% TFA in Milli-Q H₂O, and then with 100 μL of 0.2% TFA in Milli-Q H₂O, each step centrifuged at 1000 × g for 3 min. Acidified peptide samples (1% TFA) were then loaded onto the StageTips and centrifuged at 1000 × g for 3 min. The tips were washed sequentially with 100 μL of 0.2% TFA in Milli-Q H₂O and 100 μL of 99% isopropanol containing 1% TFA, each centrifuged at 1000 × g for 3 min. StageTips were transferred to fresh microcentrifuge tubes, and peptides were eluted with 100 μL of 5% ammonium hydroxide in 80% ACN by centrifugation at 1000 × g for 5 min. The eluates were dried using a SpeedVac concentrator (Labconco Centrivap Acid Resistant Centrifugal Vacuum Concentrator; no heat, 45 min) and resuspended in 50 μL of 0.1% formic acid in Milli-Q H₂O.

The samples were analysed using the Thermo Fusion ETD Tribrid (Quadrupole/Ion Trap/Orbitrap) mass spectrometer coupled to a Thermo Ultimate 3000 RSLCnano UHPLC system. Peptides were loaded onto an Acclaim PepMap 100 (100 μm x 2 cm, NanoViper C18, 5 μm, 100A, Thermo-Fisher Scientific) with 2% ACN and 0.05% TFA before switching the pre-column in line with the analytical column (Aurora Elite 15 cm x 75 μm ID, 1.7 μm C18 AUR3-15075C18-XT, Ionopticks). The separation of peptides was performed at 250 nL/min using a linear ACN gradient of buffer A (0.1% (v/v) formic acid) and buffer B (0.1% (v/v) formic acid, 80% (v/v) ACN), starting at 8% buffer B to 30% over 60 min, then rising to 50% B over 10 min followed by 97.5% B for 7 min before dropping back to 8% B. The total runtime was 90 min. Data were collected on a Thermo Fusion Orbitrap (Thermo-Fisher Scientific, Waltham, MA, USA) in Data Independent Acquisition mode, with m/z 375-1500 as the MS scan range, and HCD MS/MS spectra were collected in the Orbitrap using a loop count of 20 and a loop cycle time of 3 s. Other parameters for the instrument were: MS1 scan at 60,000 resolution and a maximum injection time of 118 ms. MS2 scan was at 15,000 resolution, 400-1000 m/z scan range, HCD collision energy 30%, AGC of 400,000 and maximum injection time of 22 ms. The isolation window of the quadrupole for the precursor was 1 m/z.

Raw data-independent acquisition (DIA) files were searched against the UniProt *Mus musculus* reference proteome using DIANN 2.3.1 in a library-based approach. The precursor- and protein-group–level false discovery rates (FDR) were controlled at 1% (q ≤ 0.01); enzyme specificity was set to Trypsin/P, with up to 2 missed cleavages permitted. Carbamidomethylation of cysteine was specified as a fixed modification, oxidation of methionine, N-terminal methionine excision, and match between runs were enabled, using DIA-NN default settings. Protein quantification was performed at the protein-group level using integrated fragment ion intensities, with DIA-NN default interference correction. Downstream data analysis, including Principal Component Analysis, was performed using the DEP: Differential Enrichment analysis of Proteomics data R package built from the limma R package using the default recommended settings as follows: each protein needed to be quantified in at least 2/3 biological replicates per group; imputation was performed post-filtration and was left-censored. Morpheus^92^ was used to generate selected heatmaps for TRP channels detected across the samples. For visualisation only, these heatmaps were displayed as log_10_(intensity + 1) values to improve representation of low-abundance signals and dynamic range, whereas statistical analyses were performed on log_2_-transformed protein-group intensities. Gene Ontology cellular component (GO: CC; Supplementary Table 8) enrichment analysis, using g: Profiler^93^, was performed on the top 100 proteins (averaged across the three biological replicates) detected through each sample preparation method and presented using the ggplot2 R package.

Organ (DRG, kidney, testes) samples were processed with the same extraction chemistries, SP3/FASP workflows, LC–MS/MS acquisition method, DIA-NN search settings (1% precursor- and protein-group–level FDR), and downstream statistical workflow (DEP/limma), as described for cortex. All normalisation, filtering, and missing-value handling steps were applied identically to organ and cortex datasets to ensure comparability. Protein abundance was quantified at the protein-group level using DIA-NN, which integrates quantitative evidence across multiple peptides. Protein-group intensities were log_2_-transformed and averaged across biological replicates to obtain mean protein abundance. Quality control and quantitative assessment of proteomic data were performed using DIA-NN output matrices. The precursor-level and protein group-level intensity matrices used for downstream analyses are provided in Supplementary Tables 6 and 7, respectively. Differential protein abundance results for SN versus control and Pellet versus control comparisons are provided in Supplementary Tables 9 and 10, respectively.

Purified Xenoblot-positive control proteins for TRPA1 and TRPV1 were obtained from Alomone Labs (TRPA1 Xenoblot-positive control, Cat. No. XB-010; TRPV1 Xenoblot-positive control, Cat. No. XB-008). These *Xenopus* oocytes overexpressing rat TRP proteins were processed alongside immunoprecipitated and fractionated mouse DRG and cortical samples and were used exclusively as LC–MS/MS identification references to aid peptide-spectrum matching and target peptide identification during mass spectrometry analysis. Database searching and quantification were performed against mouse protein entries only; no rat protein entries were used for quantifying endogenous mouse proteins.

For TRPA1 and TRPV1, protein bands were manually excised around expected molecular weight markers from SDS–PAGE gels using a clean blade, taking care to minimise excess gel. Excised gel fragments were diced into ∼1 mm cubes, transferred to low-bind tubes, destained as required in 50% ACN/50 mM ABC, dehydrated in ACN, and reduced and alkylated using TCEP and IAA. Gel pieces were rehydrated with trypsin in 50 mM ABC and digested overnight at 37 °C. Peptides were recovered from the gel by sequential extraction with 50% ACN/0.1% formic acid, dried in a vacuum concentrator, and desalted on StageTips before mass spectrometry.

### Western Blotting

Fresh cortical lysate fractions were kept on ice. Total protein concentration was measured by BCA assay (Pierce™ BCA, Thermo Fisher #23225) in 96-well plates (Corning #CLS3695) against BSA standards (Thermo Fisher #23209). For each lane, 20 µg protein was brought to volume with H₂O and mixed 3:1 with NuPAGE™ LDS Sample Buffer (4X; #NP0007) supplemented to 5% β-mercaptoethanol (final 1×). Samples were heated at 70 °C for 10 min. Proteins were resolved on 4–12% Bis-Tris gels (NuPAGE™ #NW04120BOX) using 1× MES–SDS running buffer (20× stock #NP0002 diluted in Milli-Q water). 5 µL of the molecular weight markers (Precision Plus Protein WesternC Standards, Bio-Rad #161-0385; and Precision Protein StrepTactin-HRP Conjugate, Bio-Rad #161-0380) and sample (40 µL per well) were loaded. Electrophoresis was performed at 90 V for ∼75 min, adjusting for sample load and gel format. Gels were transferred to PVDF membrane (activated in 100% methanol; Sigma #179337), using a wet tank at 90 V for 60 min in the cold room with 1× Tris–glycine/20% methanol transfer buffer (from a 10× stock: 0.25 M Tris base, 2.0 M glycine; diluted to 25 mM Tris, 200 mM glycine and 20% v/v methanol). Membranes were blocked 1 h at RT on a rocker in 5% (w/v) non-fat dry milk in PBS-T (1× PBS + 0.05% (v/v) Tween-20). Blocked membranes were incubated overnight at 4 °C on a rotator with primary antibodies diluted in TBS-T (containing 0.5% milk): Anti-TRPV1 (Alomone Labs ACC-030), 1:1,000, Anti-TRPA1 (Alomone Labs ACC-037), 1:1,000, Anti-GAPDH (loading control; Sigma #G9545), 1:1,000. 1× TBS-T was 20 mM Tris, 150 mM NaCl, 0.05% Tween-20; pH 7.6 (from 10×: 200 mM Tris base, 1.5 M NaCl; pH adjusted with HCl). The next day, membranes were washed in TBS-T (3 × 5 min, RT). HRP-conjugated anti-rabbit secondary (1:10,000 in PBS-T) was applied for 1 h at RT with gentle rocking, protected from light, then washed again in PBS-T (3 × 5 min, RT). Membranes were incubated for 5 min with mixed ECL reagents (SuperSignal™ West Femto Thermo Fisher #34095; used per manufacturer’s instructions) and imaged on a ChemiDoc (Bio-Rad). Exposure settings were kept within the automatic linear range, and GAPDH was used for normalisation; Blot images were processed in FIJI v2.17.0.

### Immunoprecipitation

Immunoprecipitation was performed from adult mouse cortex and pooled DRGs. Adult mice were deeply anaesthetised, and the bilateral cortex and DRGs were rapidly dissected on ice. Cortex (one hemisphere) was minced and homogenised in 500 µL ice-cold lysis buffer (50 mM Tris-HCl, pH 8.0, 300 mM NaCl, 1% NP-40, Halt™ protease inhibitors; Thermo Fisher) using either a glass Dounce (10–15 strokes) or a handheld motorised homogeniser (three 10 s pulses with cooling between bursts), then incubated on ice for 30 min with intermittent mixing. DRGs were homogenised in 500 µL lysis buffer using a handheld homogeniser with brief vigorous pulses on ice. To minimise loss of low-abundance TRP channels to the insoluble fraction, lysates were used directly for immunoprecipitation without centrifugation. Protein A Magnetic Beads (ab214286, Abcam) were equilibrated by gentle mixing and washed 3× with 500 µL wash buffer (lysis buffer without inhibitors). For antibody coupling, 30 µL of bead slurry was incubated with 8 µg of antibody in 200 µL of wash buffer for 1 h at room temperature with gentle rotation. The following rabbit antibodies were used: anti-TRPA1 (extracellular; ACC-037, Alomone Labs), anti-TRPV1 (VR1; ACC-030, Alomone Labs), and rabbit IgG isotype control (RIC-001, Alomone Labs). Antibody-bound beads were washed twice and incubated overnight at 4 °C with 200–500 µL tissue lysate with gentle rotation. Beads were collected on a magnetic rack, unbound lysate was removed, and beads were washed 3–5× with 500 µL high-salt wash buffer (50 mM Tris-HCl, pH 8.0, 300 mM NaCl, 1% NP-40). Bound proteins were eluted by resuspending the washed beads in 50 µL 2× LDS sample buffer. After magnetic separation, eluates were transferred to clean tubes, divided into aliquots, and stored at −80 °C. For Western blotting, an aliquot was reduced with β-mercaptoethanol and boiled at 95 °C for 5 min immediately before SDS–PAGE; the remaining aliquot was processed for mass spectrometry.

### Immunofluorescence

For TRPA1 and TRPV1 immunofluorescence, mice were perfused with ice-cold artificial cerebrospinal fluid (aCSF), and brains were immersion-fixed in 4% paraformaldehyde in 0.1 M phosphate buffer (pH 7.4) at 4 °C overnight. Fixed tissue was cryoprotected in 30% sucrose in PBS until fully submerged, then sectioned at 100 µm using a vibroslicer (Campden Instruments, MA752) in PBS. Free-floating sections were heat-mediated antigen-retrieved in 10 mM sodium citrate buffer (pH 6.0) containing 0.05% Tween-20 at 90 °C for 30 min, then cooled to room temperature and equilibrated in PBS. Sections were permeabilised in 0.1% Triton X-100 in PBS for 10 min and blocked for 2 h at room temperature in PBS containing 3% bovine serum albumin (BSA) and 0.1% Triton X-100. For the detection of TRPA1 and TRPV1, two independent immunolabelling approaches were used. In the first approach, sections were incubated overnight at 4 °C with primary antibodies against TRPA1 (Alomone Labs, ACC-037) or TRPV1 (Alomone Labs, ACC-030), diluted 1:200 in blocking buffer. Following washing, sections were incubated for 2 h at room temperature with fluorophore-conjugated secondary antibodies: donkey anti-rabbit IgG (Alexa Fluor 488; Abcam, ab150061) or donkey anti-mouse IgG (Alexa Fluor 647; Abcam, ab150111), diluted 1:500 in blocking buffer. In a second, independent experiment, a directly conjugated antibody (Alomone Labs, ACC-037-F; FITC-conjugated) for TRPA1 was used at 1:200 under identical conditions, without secondary antibody incubation. Sections were washed three times in PBS containing 0.025% Tween-20 between incubation steps. Nuclei were counterstained with DAPI (1:5000 in PBS, 10 min), followed by PBS washes. Sections were mounted using Immuno-Mount (Sigma-Aldrich, F6182). Images were acquired on a Nikon Eclipse Ni confocal microscope as z-stacks with a 2 µm step size and processed in FIJI (v2.17.0).

### Statistics

*“n”* denotes independent biological replicates (individual mice). Technical replicates in qPCR (duplicates) and repeated LC–MS/MS injections were averaged and were not treated as independent observations for statistical inference. Cross-platform concordance was assessed using two-sided Pearson and Spearman correlations on log_10_-transformed expression values where indicated. Proteins were considered reproducibly detected if quantified in at least 2 of 3 biological replicates within a given extraction workflow. No samples were excluded unless they failed predefined quality-control criteria. All analyses were performed using the software versions specified above, and the code used for data processing and figure generation is available in the repository.

## Acknowledgments

This research was undertaken with the assistance of resources from the National Computational Infrastructure (NCI Australia), an NCRIS-enabled capability supported by the Australian Government. We acknowledge the contributions of Michael Devoy, Dr Fei-Ju Li and Dr Harpreet Vohra, Specialists and Manager from the Cytometry, Histology and Spatial Multiomics (CHASM) Facility at JCSMR, ANU. We acknowledge the training and support from Dr Peter Milburn and Dr Claudia Yan of the Biomolecular Resource Facility (BRF) at JCSMR, ANU. We thank Mrs Anithahini Jeyasingham for training, and the Joint Mass Spectrometry Facility (JMSF) at the Research School of Chemistry and the Research School of Biology, in partnership with CSIRO Black Mountain Site, for instrument access. We thank Professor Brett Graham (The University of Newcastle) for providing the TRPV1-Cre-eYFP mouse line. Finally, we sincerely thank the Australian Phenomics Facility (APF) and its staff, especially Ms Barbara Burke, for animal care and husbandry.

## Author Contributions

M.B. and K.S.K. performed all experiments, led data analysis, prepared the data and figures for publication, and drafted the initial manuscript. K.S.K., M.B., R.H., and E.K. contributed to the overall study design, data interpretation, and manuscript editing. A.S. and R.H. contributed to transcriptomic sequencing experiments, developed basic transcriptomic analysis code, and contributed to the associated computational analyses. N.V. and J.S. contributed to LC–MS/MS experiments, developed proteomics workflows, and contributed to proteomics analyses. All authors have read and agreed to the published version of the manuscript.

## Institutional Review Board Statement

The animal study protocol was approved by the Animal Experimentation Ethics Committee of The Australian National University (protocol code A/2023/304: The role of calcium in physiology and pathology of rodent cortex).

## Data Availability

RNA sequencing data generated in this study have been deposited in the NCBI Gene Expression Omnibus (GEO) under accession number GSE315051 and will be publicly available upon publication. Raw FASTQ files are accessible via the associated Sequence Read Archive (SRA) records, and processed expression matrices are available in GEO. Mass spectrometry proteomics data have been deposited to the ProteomeXchange Consortium via the PRIDE partner repository with the dataset identifier PXD074939.

## Code Availability

Analysis codes to reproduce the results are available at: https://github.com/mmbilal425/TRP-channel-hierarchy-mouse-cortex.

## References

1 Montell, C. & Rubin, G. M. Molecular characterization of the Drosophila trp locus: a putative integral membrane protein required for phototransduction. Neuron 2, 1313–1323 (1989). 10.1016/0896-6273(89)90069-x

2 Nilius, B. & Owsianik, G. The transient receptor potential family of ion channels. Genome Biol 12, 218 (2011). 10.1186/gb-2011-12-3-218

3 Zhang, M. et al. TRP (transient receptor potential) ion channel family: structures, biological functions and therapeutic interventions for diseases. Signal Transduction and Targeted Therapy 8, 261 (2023). 10.1038/s41392-023-01464-x

4 Damann, N., Voets, T. & Nilius, B. TRPs in our senses. Curr Biol 18, R880–889 (2008). 10.1016/j.cub.2008.07.063

5 Wong, F. et al. Proper function of the Drosophila trp gene product during pupal development is important for normal visual transduction in the adult. Neuron 3, 81–94 (1989). 10.1016/0896-6273(89)90117-7

6 Zheng, J. Molecular mechanism of TRP channels. Compr Physiol 3, 221–242 (2013). 10.1002/cphy.c120001

7 Venkatachalam, K. & Montell, C. TRP channels. Annu Rev Biochem 76, 387–417 (2007). 10.1146/annurev.biochem.75.103004.142819

8 Moran, M. M., McAlexander, M. A., Bíró, T. & Szallasi, A. Transient receptor potential channels as therapeutic targets. Nature Reviews Drug Discovery 10, 601–620 (2011). 10.1038/nrd3456

9 Morelli, M. B., Amantini, C., Liberati, S., Santoni, M. & Nabissi, M. TRP channels: new potential therapeutic approaches in CNS neuropathies. CNS Neurol Disord Drug Targets 12, 274–293 (2013). 10.2174/18715273113129990056

10 Kheradpezhouh, E., Tang, M. F., Mattingley, J. B. & Arabzadeh, E. Enhanced Sensory Coding in Mouse Vibrissal and Visual Cortex through TRPA1. Cell Reports 32 (2020). 10.1016/j.celrep.2020.107935

11 Koivisto, A. P., Belvisi, M. G., Gaudet, R. & Szallasi, A. Advances in TRP channel drug discovery: from target validation to clinical studies. Nat Rev Drug Discov 21, 41–59 (2022). 10.1038/s41573-021-00268-4

12 Fallah, H. P. et al. A Review on the Role of TRP Channels and Their Potential as Drug Targets_An Insight Into the TRP Channel Drug Discovery Methodologies. Front Pharmacol 13, 914499 (2022). 10.3389/fphar.2022.914499

13 Clapham, D. E. TRP channels as cellular sensors. Nature 426, 517–524 (2003). 10.1038/nature02196

14 Lee, K. et al. Functional Importance of Transient Receptor Potential (TRP) Channels in Neurological Disorders. Front Cell Dev Biol 9, 611773 (2021). 10.3389/fcell.2021.611773

15 Story, G. M. et al. ANKTM1, a TRP-like channel expressed in nociceptive neurons, is activated by cold temperatures. Cell 112, 819–829 (2003). 10.1016/s0092-8674(03)00158-2

16 Caterina, M. J. et al. The capsaicin receptor: a heat-activated ion channel in the pain pathway. Nature 389, 816–824 (1997). 10.1038/39807

17 Tominaga, M. et al. The cloned capsaicin receptor integrates multiple pain-producing stimuli. Neuron 21, 531–543 (1998). 10.1016/s0896-6273(00)80564-4

18 Julius, D. TRP channels and pain. Annu Rev Cell Dev Biol 29, 355–384 (2013). 10.1146/annurev-cellbio-101011-155833

19 Paulsen, C. E., Armache, J. P., Gao, Y., Cheng, Y. & Julius, D. Structure of the TRPA1 ion channel suggests regulatory mechanisms. Nature 520, 511–517 (2015). 10.1038/nature14367

20 Kobayashi, K. et al. Distinct expression of TRPM8, TRPA1, and TRPV1 mRNAs in rat primary afferent neurons with adelta/c-fibers and colocalization with trk receptors. J Comp Neurol 493, 596–606 (2005). 10.1002/cne.20794

21 Kheradpezhouh, E., Choy, J. M. C., Daria, V. R. & Arabzadeh, E. TRPA1 expression and its functional activation in rodent cortex. Open Biol 7 (2017). 10.1098/rsob.160314

22 Liapi, A. & Wood, J. N. Extensive co-localization and heteromultimer formation of the vanilloid receptor-like protein TRPV2 and the capsaicin receptor TRPV1 in the adult rat cerebral cortex. Eur J Neurosci 22, 825–834 (2005). 10.1111/j.1460-9568.2005.04270.x

23 Mezey, E. et al. Distribution of mRNA for vanilloid receptor subtype 1 (VR1), and VR1-like immunoreactivity, in the central nervous system of the rat and human. Proc Natl Acad Sci U S A 97, 3655–3660 (2000). 10.1073/pnas.97.7.3655

24 Hodge, R. D. et al. Conserved cell types with divergent features in human versus mouse cortex. Nature 573, 61–68 (2019). 10.1038/s41586-019-1506-7

25 Leung, S. K. et al. Full-length transcript sequencing of human and mouse cerebral cortex identifies widespread isoform diversity and alternative splicing. Cell Rep 37, 110022 (2021). 10.1016/j.celrep.2021.110022

26 Sessegolo, C. et al. Transcriptome profiling of mouse samples using nanopore sequencing of cDNA and RNA molecules. Scientific Reports 9, 14908 (2019). 10.1038/s41598-019-51470-9

27 Tasic, B. et al. Adult mouse cortical cell taxonomy revealed by single cell transcriptomics. Nature Neuroscience 19, 335–346 (2016). 10.1038/nn.4216

28 Toyama, Brandon H. et al. Identification of Long-Lived Proteins Reveals Exceptional Stability of Essential Cellular Structures. Cell 154, 971–982 (2013). 10.1016/j.cell.2013.07.037

29 Bergmann, C., Mousaei, K., Rizzoli, S. O. & Tchumatchenko, T. How energy determines spatial localisation and copy number of molecules in neurons. Nature Communications 16, 1424 (2025). 10.1038/s41467-025-56640-0

30 McShane, E. et al. Kinetic Analysis of Protein Stability Reveals Age-Dependent Degradation. Cell 167, 803–815.e821 (2016). 10.1016/j.cell.2016.09.015

31 Wartenberg, P. et al. Combining mass spectrometry and genetic labeling in mice to report TRP channel expression. MethodsX 9, 101604 (2022). 10.1016/j.mex.2021.101604

32 Whitelegge, J. P. Integral Membrane Proteins and Bilayer Proteomics. Analytical Chemistry 85, 2558–2568 (2013). 10.1021/ac303064a

33 Kollewe, A. et al. Subunit composition, molecular environment, and activation of native TRPC channels encoded by their interactomes. Neuron 110, 4162–4175.e4167 (2022). 10.1016/j.neuron.2022.09.029

34 Zhang, L. J. et al. Proteomic analysis of low-abundant integral plasma membrane proteins based on gels. Cellular and Molecular Life Sciences 63, 1790 (2006). 10.1007/s00018-006-6126-3

35 Zhao, Y., Zhang, W., Kho, Y. & Zhao, Y. Proteomic Analysis of Integral Plasma Membrane Proteins. Analytical Chemistry 76, 1817–1823 (2004). 10.1021/ac0354037

36 Jordt, S. E. et al. Mustard oils and cannabinoids excite sensory nerve fibres through the TRP channel ANKTM1. Nature 427, 260–265 (2004). 10.1038/nature02282

37 Cavanaugh, D. J. et al. Restriction of Transient Receptor Potential Vanilloid-1 to the Peptidergic Subset of Primary Afferent Neurons Follows Its Developmental Downregulation in Nonpeptidergic Neurons. The Journal of Neuroscience 31, 10119–10127 (2011). 10.1523/jneurosci.1299-11.2011

38 Wang, X. L. et al. Effects of TRPA1 activation and inhibition on TRPA1 and CGRP expression in dorsal root ganglion neurons. Neural Regen Res 14, 140–148 (2019). 10.4103/1673-5374.243719

39 Kunert-Keil, C., Bisping, F., Krüger, J. & Brinkmeier, H. Tissue-specific expression of TRP channel genes in the mouse and its variation in three different mouse strains. BMC Genomics 7, 159 (2006). 10.1186/1471-2164-7-159

40 Abiria, S. A. et al. TRPM7 senses oxidative stress to release Zn^2+^ from unique intracellular vesicles. Proceedings of the National Academy of Sciences 114, E6079–E6088 (2017). doi:10.1073/pnas.1707380114

41 Jeon, J. & Xi Zhu, M. TRPC4 Channels are a Key Player in Hippocampal Neuronal Development. Biophysical Journal 116, 538a (2019). 10.1016/j.bpj.2018.11.2893

42 Schattling, B. et al. TRPM4 cation channel mediates axonal and neuronal degeneration in experimental autoimmune encephalomyelitis and multiple sclerosis. Nat Med 18, 1805–1811 (2012). 10.1038/nm.3015

43 Jang, Y. et al. Quantitative analysis of TRP channel genes in mouse organs. Arch Pharm Res 35, 1823–1830 (2012). 10.1007/s12272-012-1016-8

44 Patowary, A. et al. Developmental isoform diversity in the human neocortex informs neuropsychiatric risk mechanisms. Science 384, eadh7688 (2024). 10.1126/science.adh7688

45 Yao, Z. et al. A high-resolution transcriptomic and spatial atlas of cell types in the whole mouse brain. Nature 624, 317–332 (2023). 10.1038/s41586-023-06812-z

46 Yao, Z. et al. A taxonomy of transcriptomic cell types across the isocortex and hippocampal formation. Cell 184, 3222–3241.e3226 (2021). 10.1016/j.cell.2021.04.021

47 Zeisel, A. et al. Molecular Architecture of the Mouse Nervous System. Cell 174, 999–1014.e1022 (2018). 10.1016/j.cell.2018.06.021

48 Rostock, C., Schrenk-Siemens, K., Pohle, J. & Siemens, J. Human vs. Mouse Nociceptors - Similarities and Differences. Neuroscience 387, 13–27 (2018). 10.1016/j.neuroscience.2017.11.047

49 Bruderer, R. et al. Optimization of Experimental Parameters in Data-Independent Mass Spectrometry Significantly Increases Depth and Reproducibility of Results. Mol Cell Proteomics 16, 2296–2309 (2017). 10.1074/mcp.RA117.000314

50 Rouwette, T., Sondermann, J., Avenali, L., Gomez-Varela, D. & Schmidt, M. Standardized Profiling of The Membrane-Enriched Proteome of Mouse Dorsal Root Ganglia (DRG) Provides Novel Insights Into Chronic Pain*. Molecular & Cellular Proteomics 15, 2152–2168 (2016). 10.1074/mcp.M116.058966

51 Schmidt, M. et al. Transcriptomic and proteomic profiling of NaV1.8-expressing mouse nociceptors. Frontiers in Molecular Neuroscience Volume 15 **-** 2022 (2022). 10.3389/fnmol.2022.1002842

52 Xiong, X. et al. Enrichment and proteomic analysis of plasma membrane from rat dorsal root ganglions. Proteome Science 7, 41 (2009). 10.1186/1477-5956-7-41

53 Startek, J. B. et al. Mouse TRPA1 function and membrane localization are modulated by direct interactions with cholesterol. eLife 8, e46084 (2019). 10.7554/eLife.46084

54 da Costa, D. S. M. et al. The involvement of the transient receptor potential A1 (TRPA1) in the maintenance of mechanical and cold hyperalgesia in persistent inflammation. PAIN 148 (2010).

55 Patil, M. J. et al. A Novel Flp Reporter Mouse Shows That TRPA1 Expression Is Largely Limited to Sensory Neuron Subsets. eneuro 10, ENEURO.0350-0323.2023 (2023). 10.1523/eneuro.0350-23.2023

56 Bhattacharjee, A., Abuammar, H. & Juhász, G. Lysosomal activity depends on TRPML1-mediated Ca(2+) release coupled to incoming vesicle fusions. J Biol Chem 300, 107911 (2024). 10.1016/j.jbc.2024.107911

57 Heo, S. et al. Identification of long-lived synaptic proteins by proteomic analysis of synaptosome protein turnover. Proc Natl Acad Sci U S A 115, E3827–e3836 (2018). 10.1073/pnas.1720956115

58 Matveeva, A. et al. Integrated analysis of transcriptomic and proteomic alterations in mouse models of ALS/FTD identify early metabolic adaptions with similarities to mitochondrial dysfunction disorders. Amyotrophic Lateral Sclerosis and Frontotemporal Degeneration 25, 135–149 (2024). 10.1080/21678421.2023.2261979

59 Sharma, K. et al. Cell type– and brain region–resolved mouse brain proteome. Nature Neuroscience 18, 1819–1831 (2015). 10.1038/nn.4160

60 Kawabata, R. et al. TRPA1 as a O2 sensor detects microenvironmental hypoxia in the mice anterior cingulate cortex. Scientific Reports 13, 2960 (2023). 10.1038/s41598-023-29140-8

61 Cui, Y., Perez, S. & Venance, L. Endocannabinoid-LTP Mediated by CB1 and TRPV1 Receptors Encodes for Limited Occurrences of Coincident Activity in Neocortex. Front Cell Neurosci 12, 182 (2018). 10.3389/fncel.2018.00182

62 Shigetomi, E., Jackson-Weaver, O., Huckstepp, R. T., O’Dell, T. J. & Khakh, B. S. TRPA1 Channels Are Regulators of Astrocyte Basal Calcium Levels and Long-Term Potentiation via Constitutive D-Serine Release. The Journal of Neuroscience 33, 10143–10153 (2013). 10.1523/jneurosci.5779-12.2013

63 Shigetomi, E., Tong, X., Kwan, K. Y., Corey, D. P. & Khakh, B. S. TRPA1 channels regulate astrocyte resting calcium and inhibitory synapse efficacy through GAT-3. Nat Neurosci 15, 70–80 (2011). 10.1038/nn.3000

64 Virk, H. S. et al. Validation of antibodies for the specific detection of human TRPA1. Scientific Reports 9, 18500 (2019). 10.1038/s41598-019-55133-7

65 Cristino, L. et al. Immunohistochemical localization of cannabinoid type 1 and vanilloid transient receptor potential vanilloid type 1 receptors in the mouse brain. Neuroscience 139, 1405–1415 (2006). 10.1016/j.neuroscience.2006.02.074

66 Giordano, C. et al. TRPV1-dependent and -independent alterations in the limbic cortex of neuropathic mice: impact on glial caspases and pain perception. Cereb Cortex 22, 2495–2518 (2012). 10.1093/cercor/bhr328

67 Iglesias, L. P. et al. TRPV1 modulation of contextual fear memory depends on stimulus intensity and endocannabinoid signalling in the dorsal hippocampus. Neuropharmacology 224, 109314 (2023). 10.1016/j.neuropharm.2022.109314

68 Marrone, M. C. et al. TRPV1 channels are critical brain inflammation detectors and neuropathic pain biomarkers in mice. Nat Commun 8, 15292 (2017). 10.1038/ncomms15292

69 Roberts, J. C., Davis, J. B. & Benham, C. D. [3H]Resiniferatoxin autoradiography in the CNS of wild-type and TRPV1 null mice defines TRPV1 (VR-1) protein distribution. Brain Res 995, 176–183 (2004). 10.1016/j.brainres.2003.10.001

70 Sanchez, J. F., Krause, J. E. & Cortright, D. N. The distribution and regulation of vanilloid receptor VR1 and VR1 5’ splice variant RNA expression in rat. Neuroscience 107, 373–381 (2001). 10.1016/s0306-4522(01)00373-6

71 Tóth, A. et al. Expression and distribution of vanilloid receptor 1 (TRPV1) in the adult rat brain. Molecular Brain Research 135, 162–168 (2005). 10.1016/j.molbrainres.2004.12.003

72 Xu, K. et al. TRPV1-mediated sonogenetic neuromodulation of motor cortex in freely moving mice. J Neural Eng 20 (2023). 10.1088/1741-2552/acbba0

73 Huang, W. X., Min, J. W., Liu, Y. Q., He, X. H. & Peng, B. W. Expression of TRPV1 in the C57BL/6 mice brain hippocampus and cortex during development. Neuroreport 25, 379–385 (2014). 10.1097/wnr.0000000000000105

74 Puente, N. et al. The transient receptor potential vanilloid-1 is localized at excitatory synapses in the mouse dentate gyrus. Brain Struct Funct 220, 1187–1194 (2015). 10.1007/s00429-014-0711-2

75 de Novellis, V. et al. The blockade of the transient receptor potential vanilloid type 1 and fatty acid amide hydrolase decreases symptoms and central sequelae in the medial prefrontal cortex of neuropathic rats. Mol Pain 7, 7 (2011). 10.1186/1744-8069-7-7

76 Jo, K. D., Lee, K. S., Lee, W. T., Hur, M. S. & Kim, H. J. Expression of transient receptor potential channels in the ependymal cells of the developing rat brain. Anat Cell Biol 46, 68–78 (2013). 10.5115/acb.2013.46.1.68

77 Lee, K.-I. et al. Role of transient receptor potential ankyrin 1 channels in Alzheimer’s disease. Journal of Neuroinflammation 13, 92 (2016). 10.1186/s12974-016-0557-z

78 Sullivan, M. N. et al. Localized TRPA1 channel Ca2+ signals stimulated by reactive oxygen species promote cerebral artery dilation. Sci Signal 8, ra2 (2015). 10.1126/scisignal.2005659

79 Thakore, P. et al. Brain endothelial cell TRPA1 channels initiate neurovascular coupling. eLife 10, e63040 (2021). 10.7554/eLife.63040

80 Andersson, D. A., Gentry, C., Moss, S. & Bevan, S. Transient receptor potential A1 is a sensory receptor for multiple products of oxidative stress. J Neurosci 28, 2485–2494 (2008). 10.1523/jneurosci.5369-07.2008

81 McNamara, C. R. et al. TRPA1 mediates formalin-induced pain. Proceedings of the National Academy of Sciences 104, 13525–13530 (2007). doi:10.1073/pnas.0705924104

82 Nagata, K., Duggan, A., Kumar, G. & García-Añoveros, J. Nociceptor and hair cell transducer properties of TRPA1, a channel for pain and hearing. J Neurosci 25, 4052–4061 (2005). 10.1523/jneurosci.0013-05.2005

83 Günaydın, C., Arslan, G. & Bilge, S. S. Proconvulsant effect of trans-cinnamaldehyde in pentylenetetrazole-induced kindling model of epilepsy: The role of TRPA1 channels. Neuroscience Letters 721, 134823 (2020). 10.1016/j.neulet.2020.134823

84 Heydari, F. S., Gorji Valokola, M., Mehri, S., Abnous, K. & Roohbakhsh, A. The blockade of transient receptor potential ankyrin 1 (TRPA1) protects against PTZ-induced seizure. Metabolic Brain Disease 38, 621–630 (2023). 10.1007/s11011-022-01123-0

85 Bautista, D. M., Pellegrino, M. & Tsunozaki, M. TRPA1: A gatekeeper for inflammation. Annu Rev Physiol 75, 181–200 (2013). 10.1146/annurev-physiol-030212-183811

86 Paumier, A. et al. Astrocyte–neuron interplay is critical for Alzheimer’s disease pathogenesis and is rescued by TRPA1 channel blockade. Brain 145, 388–405 (2021). 10.1093/brain/awab281

87 Yang, X.-J. et al. Inhibition of TRPA1 Attenuates Oxidative Stress-induced Damage After Traumatic Brain Injury via the ERK/AKT Signaling Pathway. Neuroscience 494, 51–68 (2022). 10.1016/j.neuroscience.2022.02.003

88 Sleigh, J. N., Weir, G. A. & Schiavo, G. A simple, step-by-step dissection protocol for the rapid isolation of mouse dorsal root ganglia. BMC Res Notes 9, 82 (2016). 10.1186/s13104-016-1915-8

89 Kongpracha, P. et al. Simple But Efficacious Enrichment of Integral Membrane Proteins and Their Interactions for In-Depth Membrane Proteomics. Molecular & Cellular Proteomics 21 (2022). 10.1016/j.mcpro.2022.100206

90 Wiśniewski, J. R., Zougman, A., Nagaraj, N. & Mann, M. Universal sample preparation method for proteome analysis. Nature Methods 6, 359–362 (2009). 10.1038/nmeth.1322

91 Hughes, C. S. et al. Single-pot, solid-phase-enhanced sample preparation for proteomics experiments. Nat Protoc 14, 68–85 (2019). 10.1038/s41596-018-0082-x

92 Morpheus, B. I. Morpheus: Versatile matrix visualization and analysis software, https://software.broadinstitute.org/morpheus (2025).

93 Raudvere, U. et al. g:Profiler: a web server for functional enrichment analysis and conversions of gene lists (2019 update). Nucleic Acids Research 47, W191–W198 (2019). 10.1093/nar/gkz369

