## Supplementary material for "Integrated transcriptomics and proteomics define the TRP channel hierarchy in mouse cortex": Suplementary Figure

#### Supplementary Figure 1

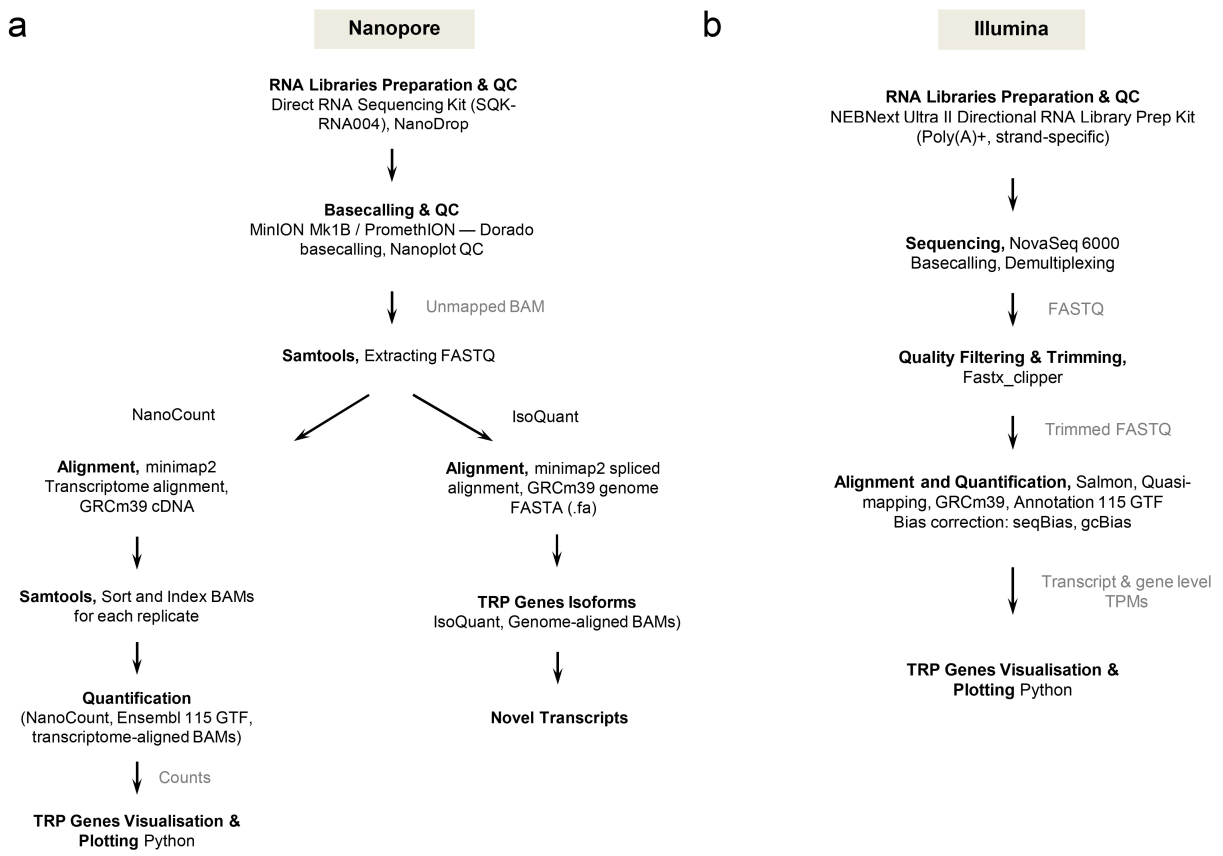


**Supplementary Figure 1: RNA-sequencing workflows for TRP profiling in adult mouse cortex.** Analysis workflow. **(a)** Nanopore direct RNA-seq libraries were basecalled (Dorado), quality controlled, aligned to GRCm39, and quantified at the gene level (NanoCount) with isoform discovery (IsoQuant; Ensembl r115). **(b)** Illumina poly(A)+ stranded RNA-seq libraries were quality controlled/trimmed and quantified by quasi-mapping (Salmon) against GRCm39/Ensembl r115.

### Supplementary Figure 2

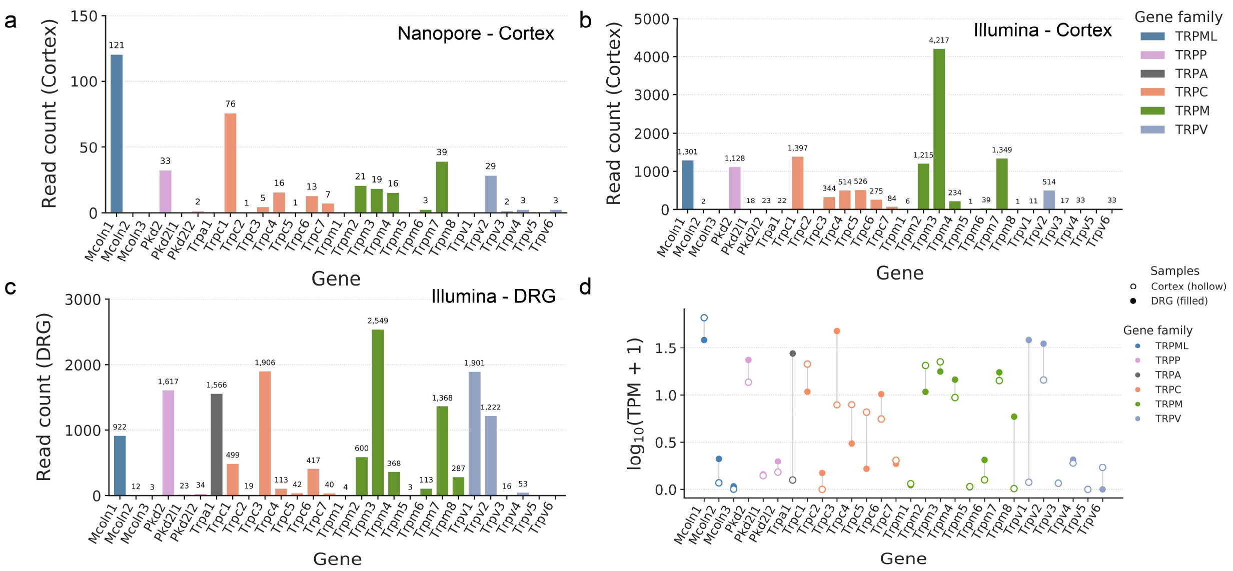


**Supplementary Figure 2: Read-count–based comparison of TRP gene expression across RNA-seq platforms and samples. (a)** Bar plot of Nanopore direct RNA-seq read counts for TRP genes in adult mouse cortex, derived from merged cortical BAM files (mean of *n* = 3 biological replicates). Genes are grouped by TRP subfamily. The relative read-count distribution is consistent with the TPM-based ranking shown in Fig. 1B. **(b)** Illumina short-read RNA-seq read counts for adult mouse cortex (mean of *n* = 3 biological replicates), showing differential representation of TRP genes in cortex. **(c)** Illumina short-read RNA-seq read counts for dorsal root ganglia (DRG; pooled sample), showing increased representation of sensory-associated TRP genes relative to cortex. **(d)** Per-gene $\log_{10}$(TPM + 1) values derived from Illumina RNA-seq for cortex (hollow symbols) and DRG (filled symbols), with lines connecting tissue pairs, highlighting divergent cortical and sensory-enriched TRP transcript profiles. Colours denote TRP subfamilies.

### Supplementary Figure 3

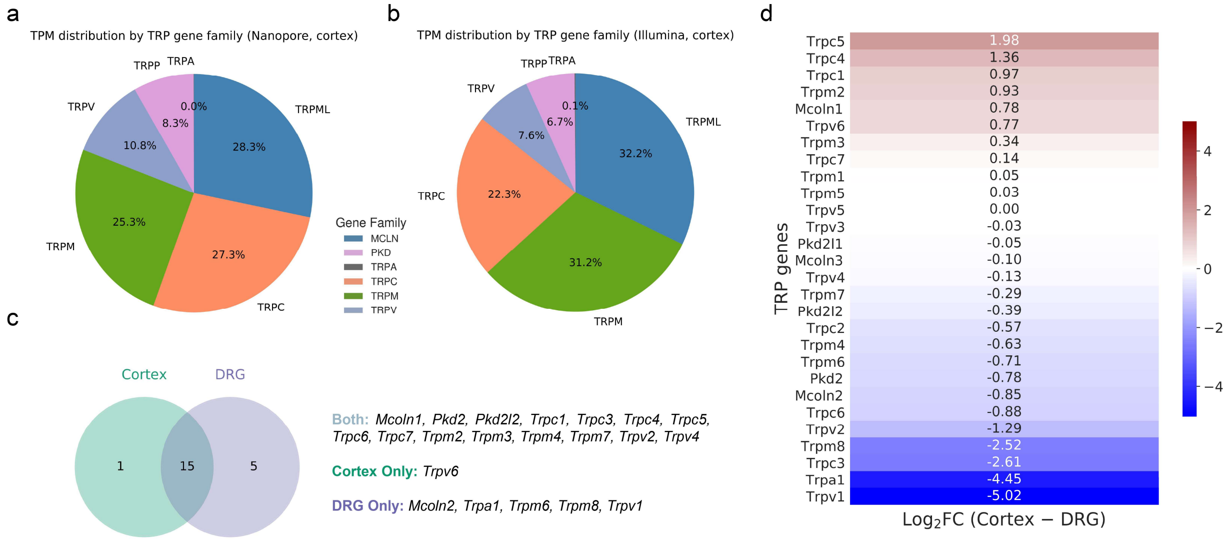


**Supplementary Figure 3: Distribution of TRP gene families across cortex and dorsal root ganglia. (a)** Pie chart showing the proportional contribution of TRP subfamilies to the cortical TRP transcript pool based on Nanopore direct RNA-seq (mean TPM across $n = 3$ biological replicates). **(b)** Pie chart of cortical TRP subfamily contributions derived from Illumina short-read RNA-seq (mean TPM across $n = 3$ biological replicates), showing a broadly similar hierarchy to Nanopore sequencing. **(c)** Venn diagram illustrating overlap of TRP genes with mean TPM $\geq 0.5$ in cortex and DRG (Illumina RNA-seq). **(d)** Heatmap of $\log_2$(TPM + 1) fold-change (cortex − DRG), highlighting DRG-enriched sensory TRP channels (*Trpa1, Trpv1, Trpc3* and *Trpm8*) relative to cortical TRP transcripts.

**Isoform reconstruction identifies a limited set of candidate novel TRP splice variants in adult cortex**: We examined whether alternative splicing contributes to TRP transcript diversity in adult mouse cortex. Although large-scale cortical transcriptomic resources catalogue extensive isoform complexity[1,2] Gene–family–resolved insights into TRP channels remain limited. Using Oxford Nanopore direct RNA sequencing, analysed with IsoQuant against the GRCm39/Ensembl release 115 annotation, we confirmed that cortical TRP expression is dominated by a small number of annotated (full-splice-match) transcripts (Supplementary Fig. 4a). IsoQuant additionally identified seven non-redundant TRP transcript models, not represented as complete transcript models in Ensembl r115, that were reproducibly detected in at least 2 of 3 biological replicates (Supplementary Fig. 4b). These comprised five novel-in-catalog (NIC) isoforms (*Mcoln1* transcript42.8.nic, *Pkd2l2* transcript1448.18.nic, *Trpv2* transcript4416.11.nic, *Trpc4* transcript1443.3.nic, and *Trpv6* transcript1584.6.nic) and two novel-not-in-catalog (NNC) isoforms (*Trpc7* transcript2626.13.nnic and *Pkd2* transcript4876.5.nnic).

All candidate novel isoforms were reproducibly detected but expressed at low abundance (mean TPM ~4; range ~1.7–7.5), indicating that alternative splicing contributes a modest additional layer of transcript diversity on top of the generally low basal expression of TRP channels in adult cortex. Structural inspection suggested exon skipping and alternative splice-site usage across multiple TRP loci, nominating these transcripts as candidates for future validation and functional investigation. For example, our isoform-resolved calls for *Trpm7* and *Trpc1* raise testable hypotheses regarding isoform-specific gating and trafficking; while the peptide-level support for TRPV2, TRPM3/7 prioritises these as tractable targets for functional studies[3].

### Supplementary Figure 4

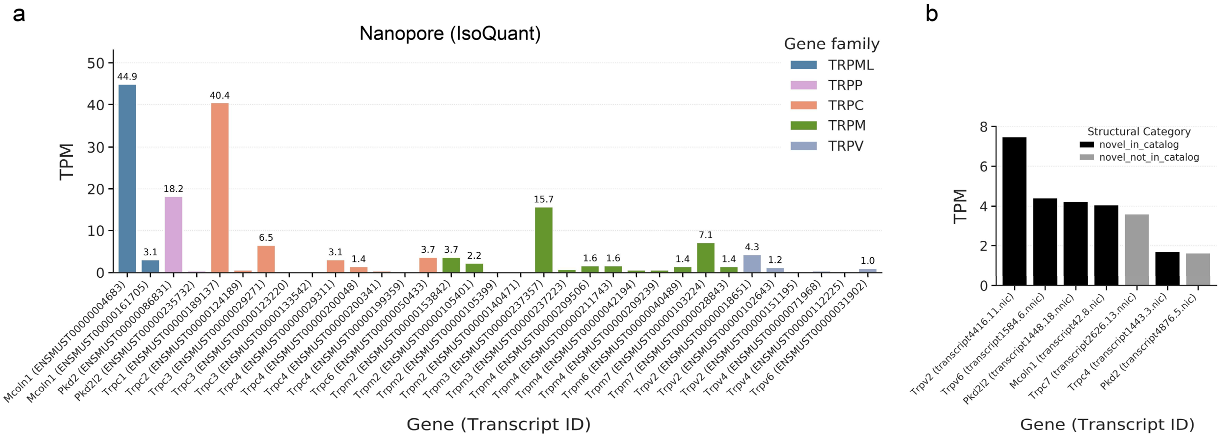


**Supplementary Figure 4: Isoform-level analysis of TRP transcripts in adult mouse cortex using Nanopore long reads. (a)** Annotated (known) TRP transcript models detected in adult cortex by IsoQuant, shown as mean TPM across $n = 3$ biological replicates; colours denote TRP subfamilies. **(b)** Candidate novel TRP transcript models detected by IsoQuant and classified by structural category as novel_in_catalog (NIC, black) or novel_not_in_catalog (NNC, grey). Bars show mean TPM for isoforms reproducibly detected across biological replicates. Reference annotation: GRCm39/Ensembl release 115. X-axis labels indicate gene symbol and IsoQuant transcript ID.

#### Supplementary Figure 5

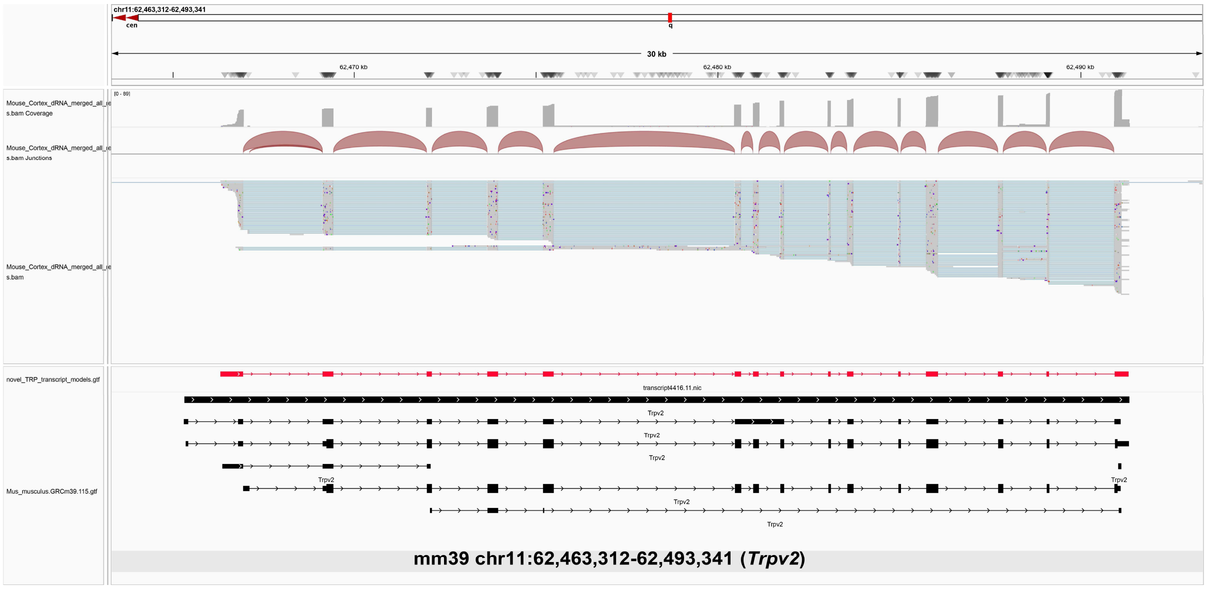


**Supplementary Figure 5: Representative IGV view of a *Trpv2* novel_in_catalog transcript model in adult mouse cortex.** Genome browser view of the *Trpv2* locus (mm39, chr11) showing merged ONT direct-RNA alignments from three biological replicates. The red transcript corresponds to the IsoQuant-derived *novel_in_catalog* model (*transcript4416.11.nic*), and black models represent Ensembl GRCm39 release 115 annotations. Junction-spanning long reads support the exon-exon boundaries of the IsoQuant model, including usage of the extended 5′ exon relative to annotated transcripts. This example illustrates how long-read data were used to inspect candidate TRP isoforms identified by IsoQuant.

**Epitranscriptomic features of cortical TRP transcripts**: We profiled $N^6$-methyladenosine ($m^6A$) RNA modification patterns on TRP transcripts using Nanopore direct RNA data, a modification implicated in regulating mRNA stability and translational efficiency[4]. Across TRP loci, $m^6A$ sites spanned a broad range of estimated stoichiometries, with most sites detected at low-to-intermediate levels and a subset approaching near-complete modification (Supplementary Fig. 6a). Motif analysis showed enrichment of the canonical DRACH consensus among high-confidence modified sites (Supplementary Fig. 6b). TRP transcripts, particularly *Trpm3*, harboured the highest number of modified positions; *Trpc3–6, Mcoln1*, *Trpm7*, *Trpv6* and *Trpm6* also exhibit enriched $m^6A$ signals (Supplementary Fig. 6a-c). This pattern aligns with our prior epitranscriptomic profiling of the adult brain[5], and suggests that post-transcriptional control may contribute to shaping TRP transcript fate. However, translation efficiency and ribosome engagement were not directly measured here, and methylation burden alone cannot predict steady-state peptide detectability. Translational output is shaped by site-specific $m^6A$ deposition, reader-protein interactions and subcellular localisation and ribosomal engagement dynamics[5-7]. More broadly, pronounced isoform heterogeneity (Supplementary Fig. 4a,b and 5) raises the possibility that subsets of TRP transcripts are translationally repressed, inefficiently trafficked or yield unstable protein products[8]. Collectively, these data indicate that cortical TRP transcripts carry substantial $m^6A$ modification, particularly within TRPM/TRPC family members, providing a framework for future studies of epitranscriptomic regulation across TRP loci.

### Supplementary Figure 6

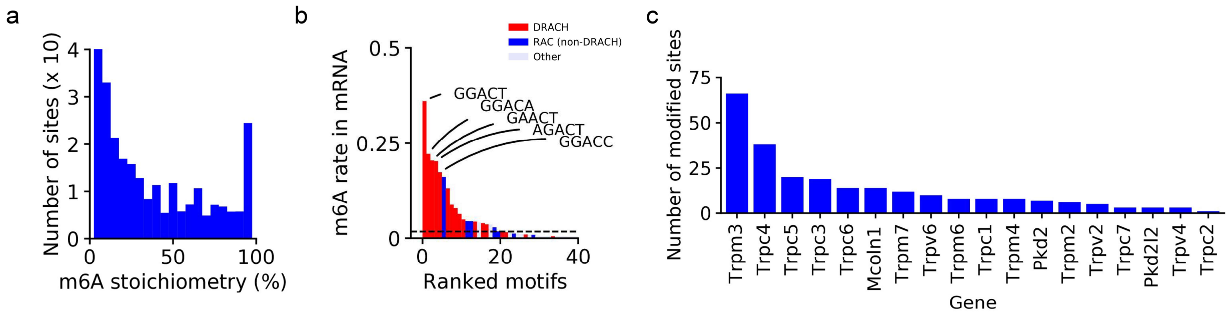


**Supplementary Figure 6: Epitranscriptomic features of TRP transcripts in adult mouse cortex.** **(a)** Distribution of estimated $m^6A$ stoichiometry across TRP-associated mRNA sites, showing predominantly low-to-intermediate modification with a subset of highly modified sites. **(b)** Ranked motif analysis for modified sites, highlighting enrichment of the canonical DRACH consensus (red) relative to RAC (non-DRACH) motifs (blue) and other sequences (grey). **(c)** Number of modified sites per TRP gene, showing the highest site counts in *Trpm3,* followed by *Trpc4–6* and *Mcoln1*.

#### Supplementary Figure 7

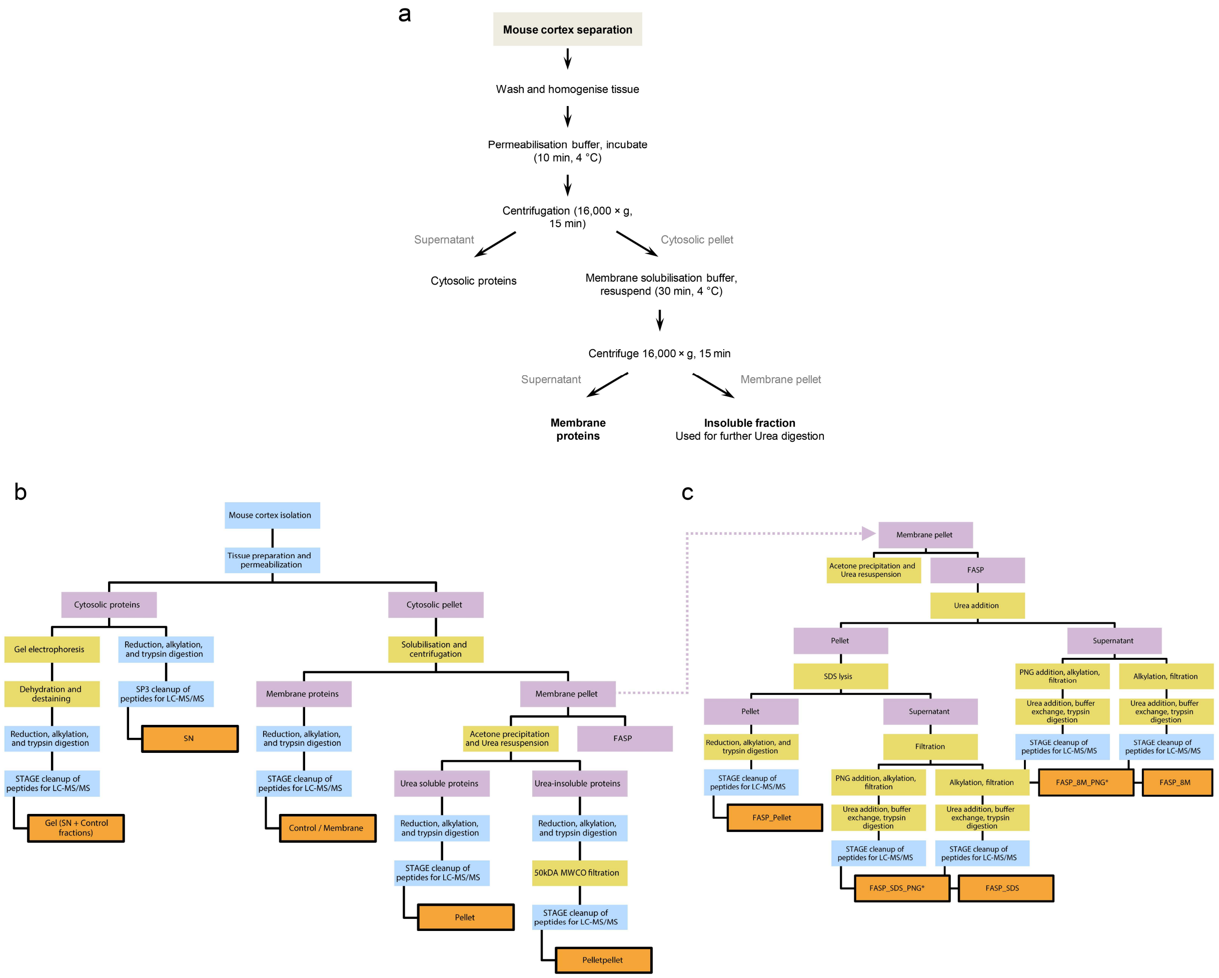


**Supplementary Figure 7. Extraction of cytosolic, membrane, urea-soluble and urea-insoluble protein fractions for LC–MS/MS. (a)** Schematic overview of the main protein extraction workflow using a Mem-PER kit–based detergent fractionation approach to separate cytosolic (supernatant) and membrane-enriched (pellet) fractions from adult mouse cortex. Following the initial extraction, the residual membrane pellet was retained for further membrane-focused processing before LC–MS/MS. Extracted proteins were reduced, alkylated, digested with trypsin, cleaned up using SP3, and analysed by LC–MS/MS. **(b, c)** Flowchart illustrating downstream processing of cytosolic, membrane, urea-soluble, and urea-insoluble fractions using alternative extraction chemistries and preparation strategies for LC–MS/MS. These included urea- and detergent-based solubilisation, acetone precipitation, molecular-weight cut-off filtration, FASP, and StageTip peptide cleanup, with terminology reflecting the corresponding sample types and preparation routes used throughout the proteomic analyses. In-gel digestion samples were generated from both cytosolic (SN) and membrane (Control) fractions following SDS–PAGE separation. Asterisk denotes exploratory PNGase F–treated samples generated to assess whether deglycosylation improves peptide recovery by reducing steric hindrance; these samples were not included in the final dataset shown.

#### Supplementary Figure 8

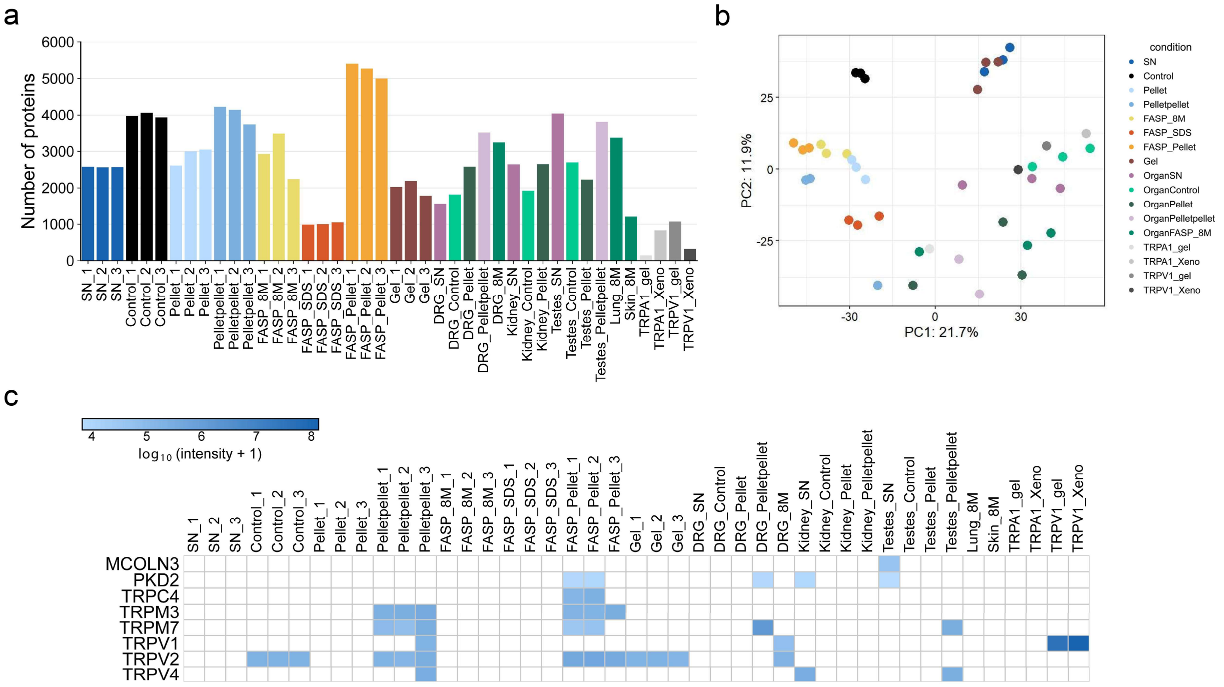


**Supplementary Figure 8. Expanded benchmarking of membrane-aware proteomics across tissues (cortex, DRG, kidney, testes). (a)** Proteins identified per sample across cortex, dorsal root ganglia (DRG), kidney, and testes, using different extraction and enrichment workflows. Membrane-enriched preparations (pellet-derived and FASP-based workflows) yielded increased proteome depth relative to supernatant (SN) and control (membrane) fractions across tissues. **(b)** Principal component analysis (PCA) of the top 500 most variable proteins across all individual LC–MS/MS samples (biological replicates; no organ-level averaging). Each point represents one sample. Replicates cluster primarily by preparation method, with membrane-enriched samples separating from SN and control preparations along PC1/PC2, indicating reproducible protocol-driven proteome differences across tissues. **(c)** Targeted TRP-focused $\log_{10}$-transformed protein intensities [$\log_{10}$(intensity + 1)] heatmap showing evidence for TRP channels across organ-specific extraction protocols. TRP recovery was strongest in FASP- and pellet-derived inputs across cortex, DRG, kidney, and testes, demonstrating that membrane-aware workflows generalise beyond cortex for the detection of low-abundance, multi-pass membrane proteins. Lung and skin 8 M urea samples were included as additional comparators, but did not show detectable TRP-channel signal in this dataset.

#### Supplementary Figure 9

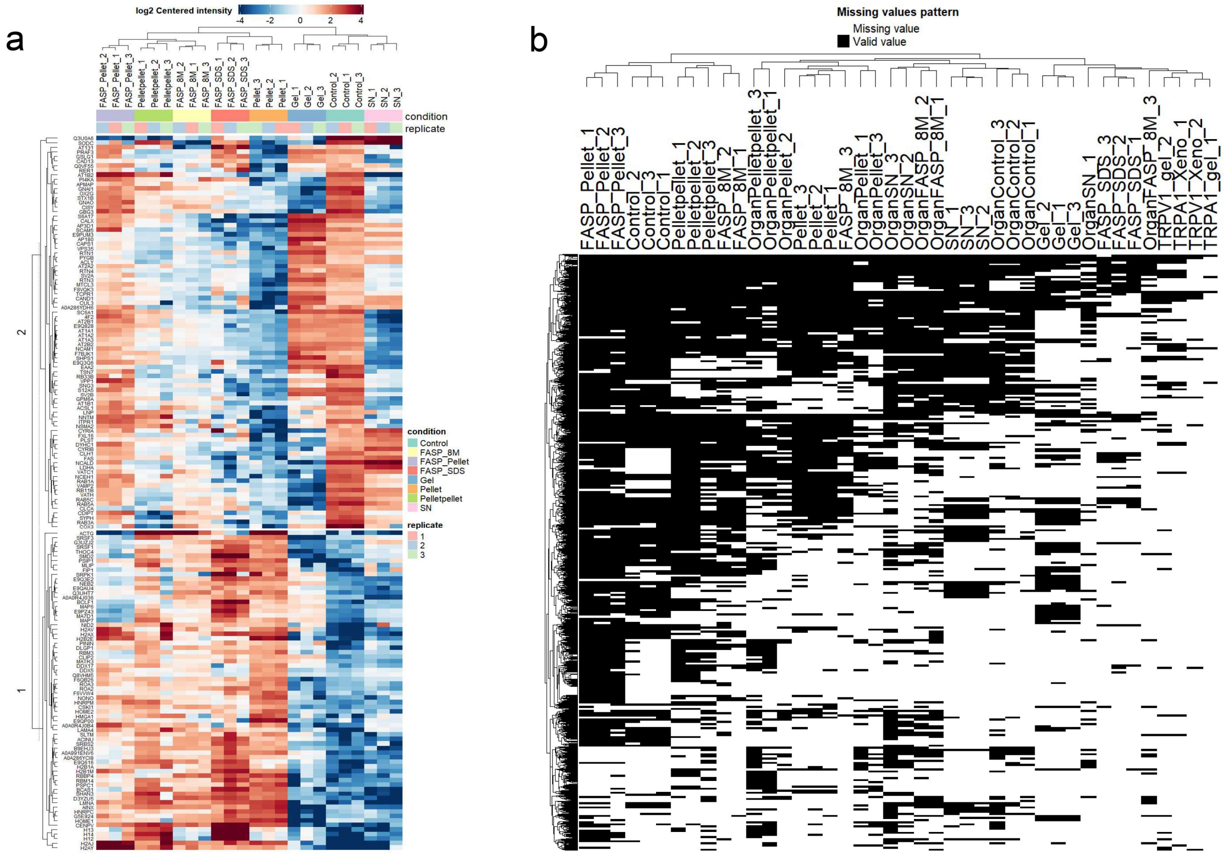


**Supplementary Figure 9. Global abundance patterns of the cortex proteome across extraction methods. (a)** Global proteomic profile across extraction methods. Heatmap of $log_2$-transformed, protein-centered intensities for the most variable proteins quantified across cortex extraction protocols. Rows = proteins; columns = biological replicates. Unsupervised hierarchical clustering groups samples by preparation method, revealing protocol-specific abundance signatures. FASP- and pellet-derived fractions segregate from supernatant (SN) and gel-based preparations, consistent with distinct proteome compositions induced by extraction chemistry. **(b)** Missing-value patterns across tissues and extraction chemistries. Binary missing-value map (black = observed; white = missing) for DRG, kidney, and testes samples processed using multiple extraction workflows. Samples cluster primarily by protocol rather than tissue, with pellet- and pelletpellet fractions exhibiting missingness patterns characteristic of membrane-enriched workflows. These profiles closely mirror the cortex missing-value structure shown in Fig. 3B, supporting protocol-driven reproducibility across organs.

#### Supplementary Figure 10

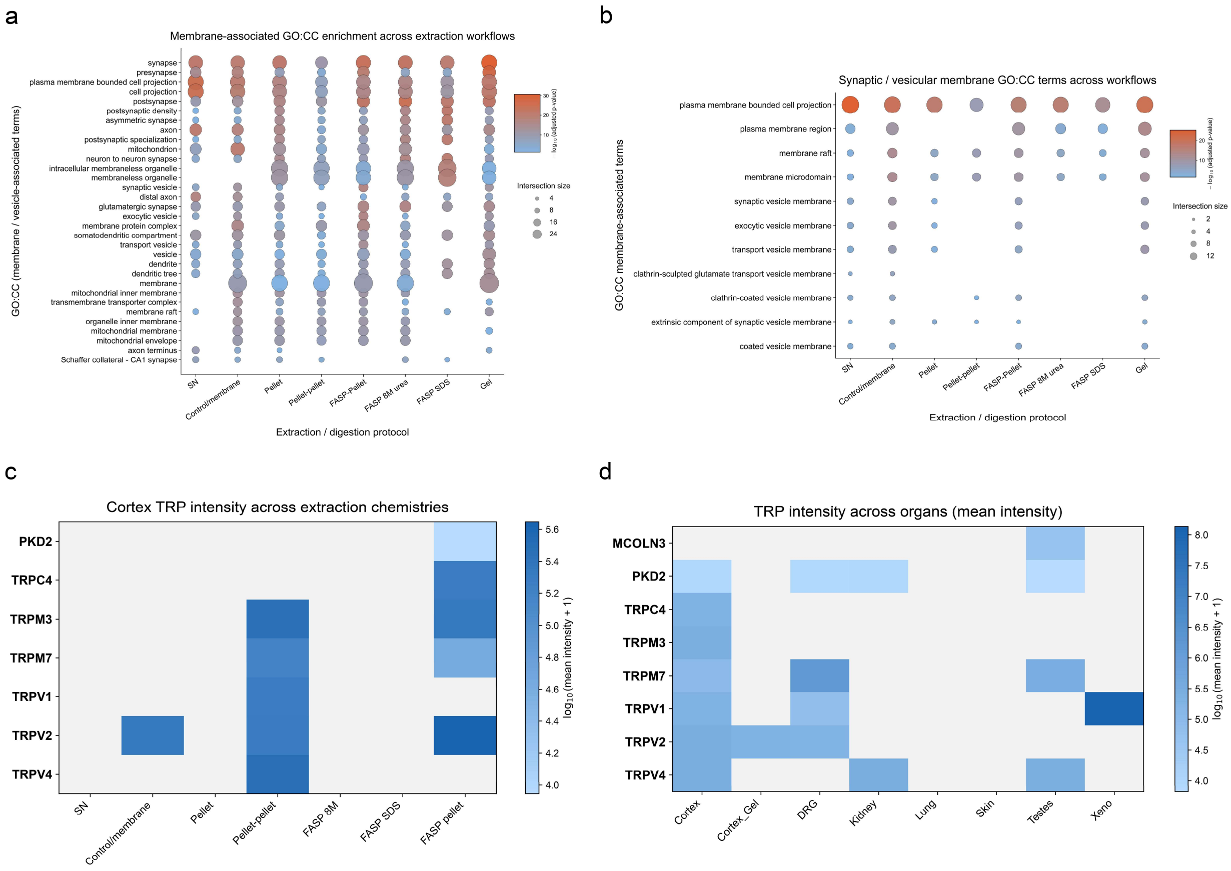


**Supplementary Figure 10. Extraction chemistry shapes membrane-associated proteome enrichment. (a)** Bubble plot of Gene Ontology Cellular Component (GO: CC) membrane- and vesicle-associated terms enriched among the top 100 proteins detected in each extraction/digestion workflows (SN/cytosolic, Control/membrane, Pellet/urea-soluble, Pelletpellet/urea-insoluble, FASP-Pellet, FASP-8M urea, FASP-SDS and Gel-excise), using the default *Mus musculus* background in g:Profiler. Columns represent extraction protocols, and rows represent GO: CC terms. Bubble size indicates intersection size (number of contributing proteins), and colour encodes enrichment significance (–$\log_{10}$ FDR-adjusted *p* value). Note: For each extraction workflow, only the top enriched membrane- and vesicle-associated GO: CC terms are shown after keyword filtering. **(b)** As in (a), focused analysis of synaptic and vesicular membrane GO: CC terms across the same extraction workflows. Bubble size indicates intersection size, and colour denotes –$\log_{10}$ FDR-adjusted *p* value. **(c)** Heatmap of mean log-transformed protein-group intensities for selected TRP channels detected in cortex across extraction chemistries. Values are shown as $\log_{10}$(mean protein-group intensity + 1). Only proteins detected in at least one condition are displayed. **(d)** Heatmap of mean log-transformed protein-group intensities for the same TRP channels across organs and control samples.

#### Supplementary Figure 11

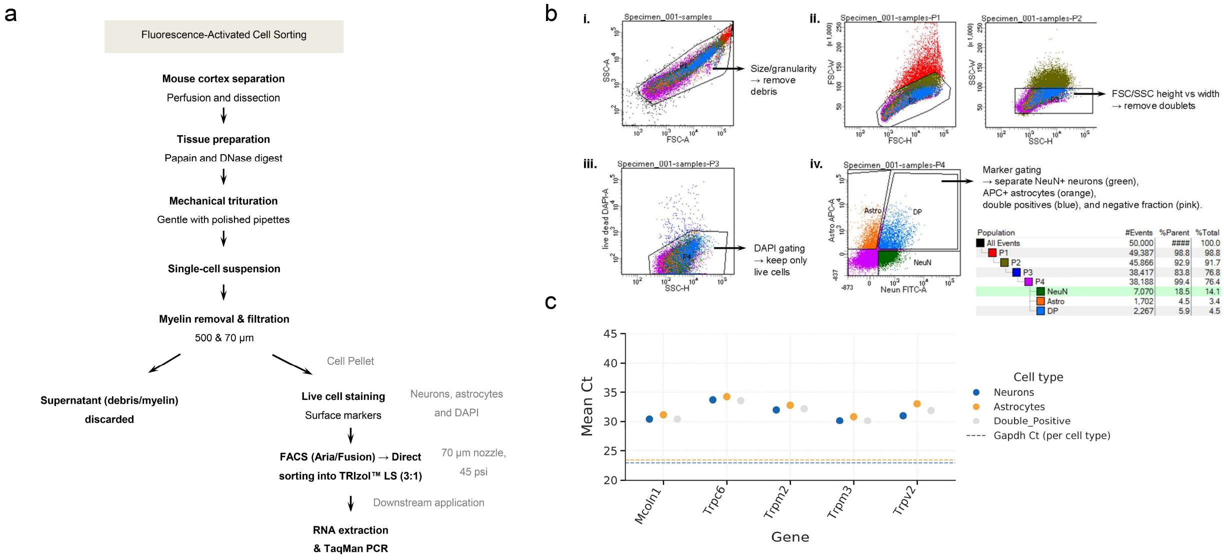


| Population | #Events | %Parent | %Total |
|---|---|---|---|
| All Events | 50,000 | #### | 100.0 |
| P1 | 49,387 | 98.8 | 98.8 |
| P2 | 45,866 | 92.9 | 91.7 |
| P3 | 38,417 | 83.8 | 76.8 |
| P4 | 38,188 | 99.4 | 76.4 |
| NeuN | 7,070 | 18.5 | 14.1 |
| Astro | 1,702 | 4.5 | 3.4 |
| DP | 2,267 | 5.9 | 4.5 |

**Supplementary Figure 11. Representative FACS gating strategy for isolating live NeuN⁺ neurons and ACSA2⁺ astrocytes from the mouse cortex. (a)** Experimental workflow for cortical dissociation, fluorescence-activated cell sorting (FACS) using NeuN and ACSA2 markers, RNA isolation, and TaqMan qPCR analysis of TRP transcripts from defined cortical cell populations. **(b)** Flow-cytometry gating: (i) FSC-A vs SSC-A to exclude debris; (ii) FSC-H vs FSC-W and SSC-H vs SSC-W to remove doublets; (iii) live/dead discrimination with DAPI to select live singlets; (iv) quadrant gating on NeuN (FITC/488) and ACSA2 (Allophycocyanin/APC) to define NeuN$^{+}$ neurons (green), ACSA2$^{+}$ astrocytes (orange), NeuN$^{+}$/ACSA2$^{+}$ double-positives (blue), and double-negatives (pink) using single-stain controls and optimised voltage/compensation. **(c)** TaqMan PCR summary by sorted fraction: *Gapdh* (~23 Ct) amplifies across all fractions; *Trpml1*, *Trpc6*, *Trpm2*, *Trpm3* and *Trpv2* amplify with Ct ~ 30-35 in neurons, astrocytes, and double-positives, indicating broad expression across cell types. Note: The NeuN$^{+}$/ACSA2$^{+}$ double-positive gate yielded housekeeping Cts comparable to pure fractions and likely contains tightly apposed neuron-astrocyte pairs or membrane aggregates rather than a true dual-marker cell population.

**Supplementary Figure 12**

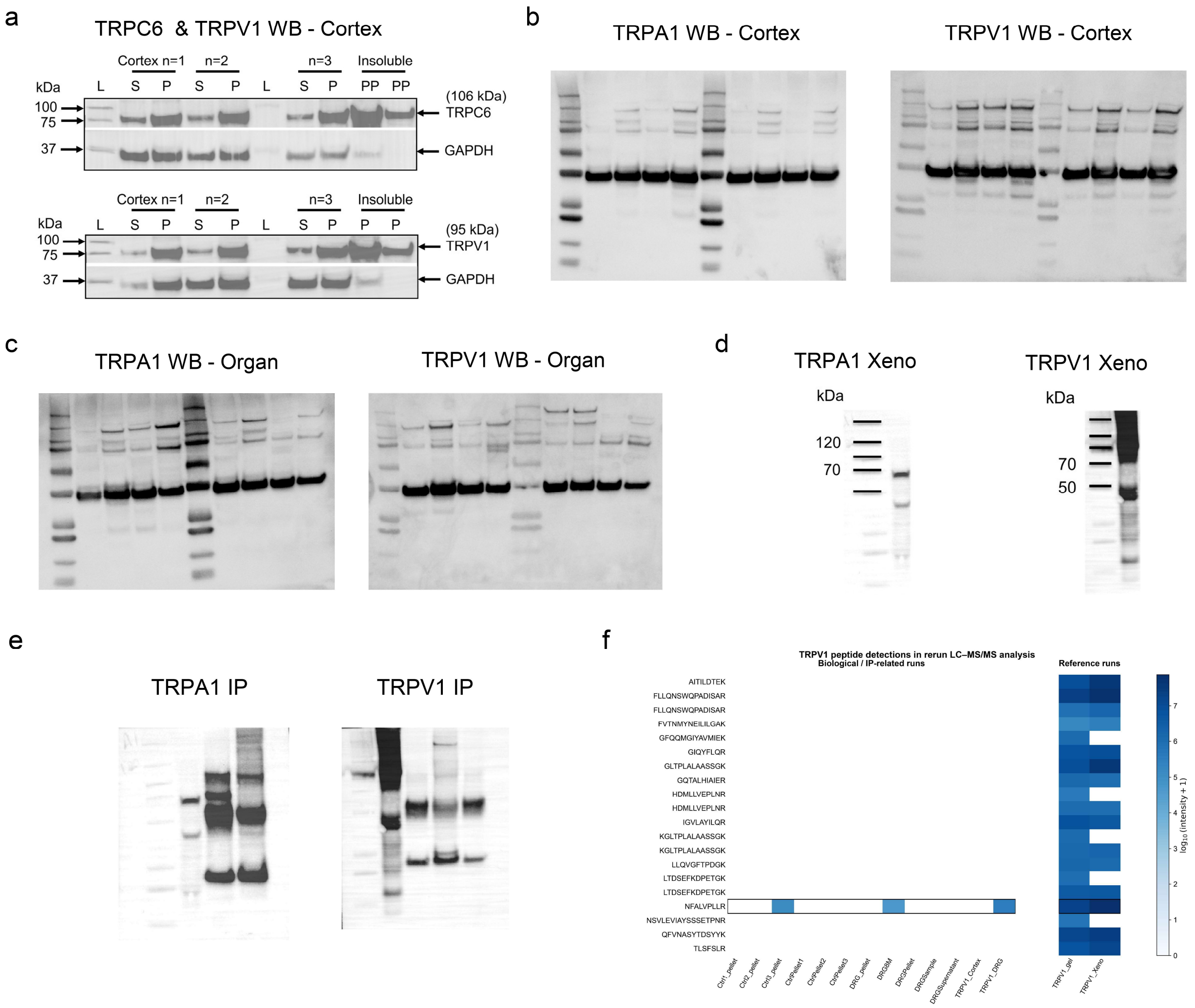


**Supplementary Figure 12. TRP protein abundance across cortex and peripheral tissues assessed by Western blot, immunoprecipitation, and LC–MS/MS peptide mapping. (a)** Representative Western blots from paired cortex fractionations showing TRPC6 (upper panel) and GAPDH (loading control) across Supernatant (S), Pellet (P), and PP (PP indicates insoluble membrane pellet fraction solubilised in SDS + 5% β-mercaptoethanol) samples (Gel 1). Representative Western blots for TRPV1 and GAPDH from the same cortex fractionation series are shown in Gel 2. Lysates were derived from three independent biological replicates (three mice; $n = 3$). All samples and controls were processed within the same experiment and run on the same gel; cropped blots were processed in parallel for each protein. **(b, c)** Unprocessed cortical and organ immunoblots correspond to Fig. 4C and 4D for TRPA1 and TRPV1. **(d)** Xenopus oocytes (Xeno) expressing recombinant rat TRPA1 and TRPV1 standards used as positive controls for LC–MS/MS peptide identification. These controls were included for reference only; due to anomalous electrophoretic mobility on SDS–PAGE, likely reflecting incomplete denaturation and/or aggregation, they were not used to infer molecular-weight concordance with endogenous mouse proteins. **(e)** Unprocessed immunoprecipitation (IP) eluates for TRPA1 and TRPV1 corresponding to Fig. 4E and 4F. Xeno control lanes shown in panel (d) were derived from the same immunoblot but are displayed separately for clarity. **(f)** Heatmap showing

TRPV1 peptide detections across LC–MS/MS reruns incorporating reference controls. Rows indicate individual detected peptides and columns indicate reruns/samples, showing peptide-level evidence for TRPV1 across TRPV1-positive preparations. One shared TRPV1 peptide was detected in a single cortex replicate and in DRG samples.

#### Supplementary Figure 13

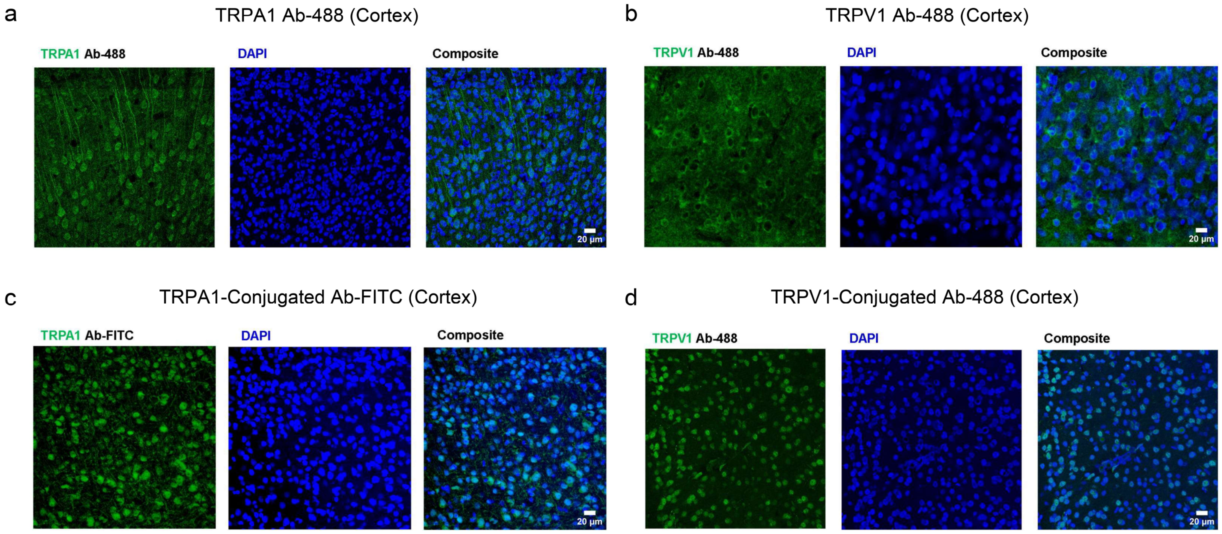


**Supplementary Figure 13. Cortical immunofluorescence staining of TRPA1 and TRPV1.** Representative confocal images from adult mouse cortex stained for TRPA1 or TRPV1 (green) with nuclear counterstain DAPI (blue). Panels **(a-d)** each show (left) target channel, (middle) nuclei (DAPI), and (right) merged image. **a,c**: TRPA1. **b,d**: TRPV1. **a,b**: Ab-488 secondary. **c**: FITC-conjugated antibody (TRPA1). **d**: Ab-488–conjugated antibody (TRPV1). Scale bars, 20 µm. Note: Immunoreactive signal alone was not interpreted as evidence of endogenous protein presence and is shown only to illustrate the limitations of antibody-based detection at the sensitivity limits of bulk tissue. We relied on peptide-level evidence as the primary criterion for protein detection.

#### Supplementary Figure 14

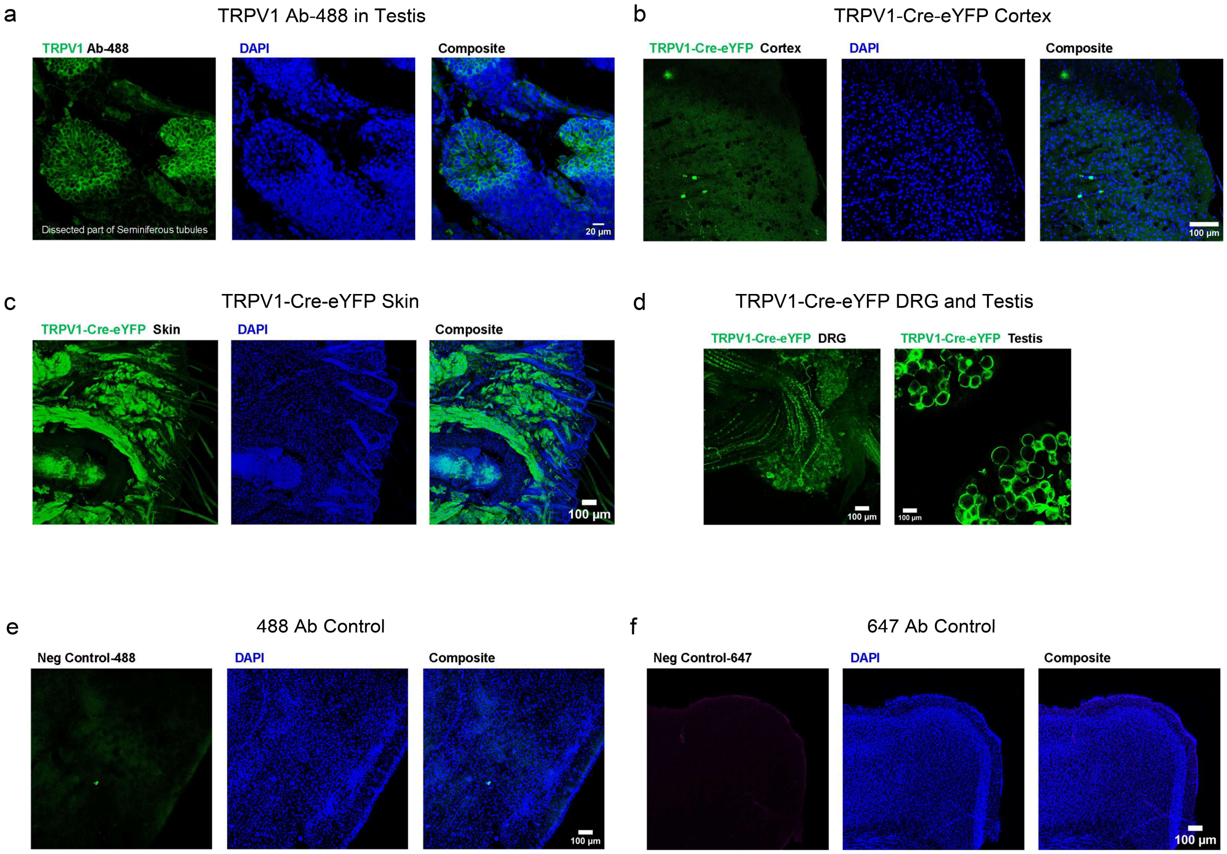


**Supplementary Figure 14. Confocal images showing immunofluorescence of TRPV1. (a)** Representative confocal images of adult mouse testis stained with a TRPV1 antibody, shown as positive control. **(b-d)** Confocal images from TRPV1-Cre-eYFP reporter mice showing eYFP signal in brain, skin, DRG, and testis, with nuclei counterstained with DAPI where indicated. **(e, f)** Negative control sections imaged in the 568- and 647-nm channels show no detectable signal, confirming labelling specificity and minimal non-specific fluorescence. Scale bars as indicated.

#### References


1 Leung, S. K. *et al.* Full-length transcript sequencing of human and mouse cerebral cortex identifies widespread isoform diversity and alternative splicing. *Cell Rep* **37**, 110022 (2021). https://doi.org:10.1016/j.celrep.2021.110022

2 Patowary, A. *et al.* Developmental isoform diversity in the human neocortex informs neuropsychiatric risk mechanisms. *Science* **384**, eadh7688 (2024). https://doi.org:10.1126/science.adh7688

3 Luo, H., Declèves, X. & Cisternino, S. Transient Receptor Potential Vanilloid in the Brain Gliovascular Unit: Prospective Targets in Therapy. *Pharmaceutics* **13**, 334 (2021).

4 Yen, Y. P. & Chen, J. A. The m(6)A epitranscriptome on neural development and degeneration. *J Biomed Sci* **28**, 40 (2021). https://doi.org:10.1186/s12929-021-00734-6

5 Sethi, A. J. *et al.* Single-molecule multimodal timing of *in vivo* mRNA synthesis. *bioRxiv*, 2025.2004.2027.650906 (2025). https://doi.org:10.1101/2025.04.27.650906

6 Meyer, K. D. *et al.* Comprehensive analysis of mRNA methylation reveals enrichment in 3' UTRs and near stop codons. *Cell* **149**, 1635-1646 (2012). https://doi.org:10.1016/j.cell.2012.05.003

7 Yoon, K. J. *et al.* Temporal Control of Mammalian Cortical Neurogenesis by m(6)A Methylation. *Cell* **171**, 877-889.e817 (2017). https://doi.org:10.1016/j.cell.2017.09.003

8 Sison, S. L. *et al.* Single-cell and isoform-specific translational profiling of the mouse brain. *Nature* (2026). https://doi.org:10.1038/s41586-026-10118-1
